# Single-cell mapping links neurogenesis and angiogenesis in aggressive breast cancer

**DOI:** 10.1101/2022.01.28.477898

**Authors:** Elisabeth Wik, Sura Aziz, Kenneth Finne, Dimitrios Kleftogiannis, Cecilie Askeland, Gøril Knutsvik, Kristi Krüger, Amalie A. Svanøe, Even Birkeland, Silje Kjølle, Benedicte Davidsen, Ingunn M. Stefansson, Heidrun Vethe, Lars A. Akslen

## Abstract

The tumor microenvironment (TME) is important for cancer growth and progression. While angiogenesis is an established hallmark of cancer, the role of nerve fibers is less studied. Here, we investigated neurogenesis and angiogenesis in breast cancer and found them to be closely associated. Single-cell based spatial mapping by imaging mass cytometry (IMC) indicated close proximity between neural and vascular structures. Subsequent validation by tissue-based markers of neurogenesis and angiogenesis, supported by proteomics and transcriptomics data of tissues and cell lines, supported a link between these processes. A consolidated neuro-angiogenic signature score was linked to high-grade breast cancer and reduced patient survival, also within the low-grade luminal tumor subgroup. Our findings support that neurogenesis and angiogenesis are related in aggressive breast cancer and might possibly improve tumor stratification and clinical management.

## Introduction

Tumor-stroma communication is important for cancer progression and is suggested as a therapeutic target. Programs of inflammation, angiogenesis, and fibroblast activation are among the tumor microenvironment (TME) characteristics of aggressive cancer (1, 2). Recently, evidence has indicated links between tumor cells, nerve elements and progressive disease (3, 4), and experimental studies suggest that cancer cells may simultaneously promote the recruitment of both sprouting axons and endothelial cells(5–9). However, studies of neural tissue in breast cancer stroma are limited (10–16), and the potential coexistence of microaxons and microvessels have, to our knowledge, not been examined in human tumor tissues.

Here, single-cell imaging mass cytometry (IMC), combined with mass spectrometry-based proteomics and transcriptomics data from different human breast cancer cohorts, were investigated for the relationship between neural and vascular processes in breast cancer TME. Our findings support the notion that neurogenesis and angiogenesis are linked in aggressive breast cancer and associated with clinical outcome. These results provide a basis for improved tumor stratification and more precise clinical management of breast cancer patients.

## Methods

### Patient series

The study cohort included women diagnosed with primary invasive breast cancer as part of the prospective and population-based Norwegian Breast Cancer Screening Program during 1996-2003 (Hordaland County, Norway; 10% of the Norwegian population), age 50–69 years at time of diagnosis (median 59 years). Detailed information about this study population and the specimen characteristics is presented in Supplementary Material.

### Imaging mass cytometry (IMC)

#### IMC breast cancer cohort

We analyzed tissues from two independent tissue collection studies (n=117). Cohort-1 consisted of tumors from the Norwegian Breast Cancer Screening Program. Cohort-2 was established in collaboration with Prof. William D. Foulkes at McGill University, Canada. We assessed both cohorts using tissue microarrays (TMAs) with 1mm cores in triplets and analyzed one core from each patient.

#### IMC antibody panel

The antibody panel consisted of 36 metal-conjugated antibodies. All antibodies were validated by IHC using a test-TMA with positive control tissues. A pilot-TMA was established to validate and calibrate the experimental protocol. Antibody hybridization was performed as described by Fluidigm with a few modifications (see Supplementary Material).

#### IMC acquisition and data pre-processing

Data from the IMC cohort (n=117) were acquired by a Helios time-of-flight mass cytometer (CyTOF) coupled to a Hyperion Imaging System (Fluidigm) and administered using the CyTOF Software (v7.0.8493; Fluidigm). The square inscribed in the circular TMA cores were laser ablated at 200 Hz at a resolution of approximately 1µm^2^. Detailed information of the cell segmentation and IMC analysis workflow can be found in Supplementary Material.

### Nerve and vessel density measurements by IHC

Sections from primary tumors and the matched axillary node with the largest metastatic focus (≥ 2 mm) were used for assessment of nerves (Neurofilament antibody, DAKO M0762), and angiogenesis markers (Factor-VIII, DAKO A0082; Ki67, DAKO 7240). Vessels from adjacent normal tissue were used as controls. For further details on immunohistochemistry and assessment of nerve- and vessel density measurements (MAD, NBD, MVD and pMVD), see Supplementary Material.

### Proteomic and mRNA datasets

In this study we have used in-house datasets: Microdissected breast cancer epithelial tissue (proteomics, n=24), Secretome from breast cancer cell lines (proteomics, n=4), Cell line lysate (mRNA, n=12), and external datasets: METABRIC (mRNA, discovery cohort, n=939; validation cohort, n=845; normal-like excluded), CCLE (mRNA, n=47) and a recently published proteomics dataset by Asleh and colleagues (17) (proteomics, n=284). For more information see Supplementary Material.

### Statistics and survival analyses

Data were analyzed using SPSS (Statistical Package for the Social Sciences), Version 28.0 (Armonk, NY, USA; IBM Corp) and R-packages via Rstudio v1.3.9. A two-sided P-value less than 0.05 was considered statistically significant. Detailed descriptions of the statistical methods can be found in Supplemental Material)

### Study approval

This study was approved by the Western Regional Committee for Medical and Health Research Ethics, REC West (REK 2014/1984). Written informed consent was not obtained from the patients, but in accordance with national ethics guidelines and procedures for retrospective studies, all participants were contacted with written information on the study and asked to respond if they objected. In total, 9 patients (1.7%) did not approve of participation.

## Results

### Single-cell mapping of breast cancer tissue by imaging mass cytometry indicates neuro-vascular proximity

To investigate the single-cell spatial distribution and potential co-localization of neural and vascular structures in breast cancer, we designed an IMC panel of 36 markers (**Supplementary Table 1**) to profile 117 samples from luminal A (n=26), luminal B (n=30), HER2 enriched (n=9) and triple-negative tumors with a basal-like phenotype (n=52). In total, we identified 413,271 single cells or equally denoted cell units (*i.e.*, 218,117 from basal-like and 195,154 from luminal-like tissues), quantifying marker expression combined with spatial information of each cell unit. Initially, unsupervised clustering was used to identify basic cell phenotypic clusters of different epithelial phenotypes, T-cells, B-cells, macrophages, endothelial cells, and stromal cells (18).

We used IMC-based images to detect areas with spatial co-localization of neural and vascular structures (**Figure 1a-b**). For each endothelial cell unit, we estimated the number of neighboring cells using a k-nearest approach, and we visualized the average distance from all cell units with high Neurofilament (NF) marker expression.

**Figure 1.**
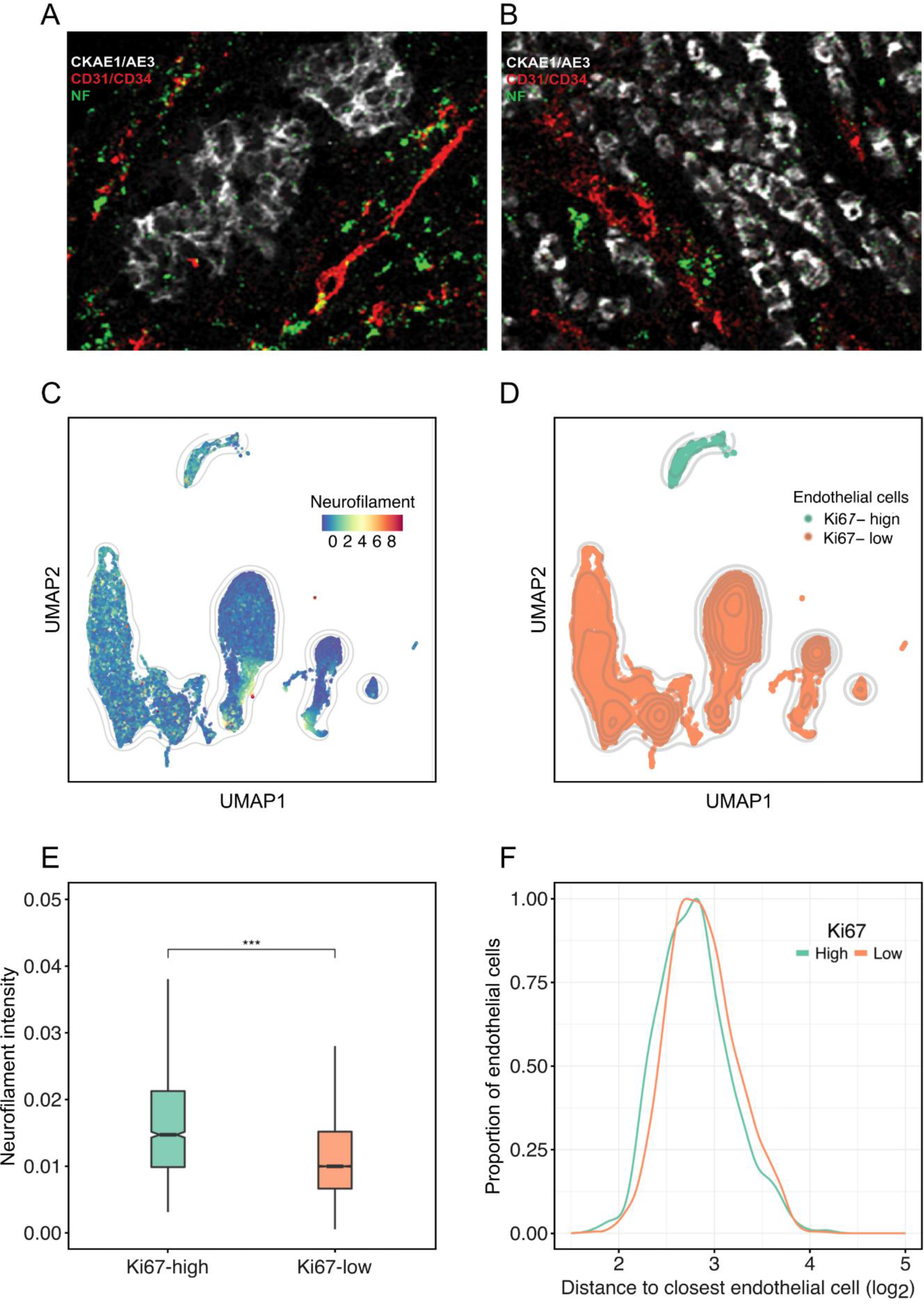
**Co-expression and spatial co-localization of neural and vascular markers and structures**. **A-B**) Representative IMC images showing co-localization of neural and vascular structures. The images are pseudo colored using the antibody intensity values of Neurofilament (green), CD31/34 (red) and the cytokeratin CKAE1/AE3 (white/grey). Focal co-expression of Neurofilament and CD31/CD34 is observed (yellow); **C-D**) UMAP projection of all endothelial cells found in the cohort of 117 patients. Single-cells are colored based on z-scored Neurofilament expression, and stratified into Ki67-high and Ki67-low groups based on Ki67 antibody intensity; **E**) Boxplot comparing neurofilament intensity in Ki67-high and Ki67-low CD31/CD34- positive endothelial cells (p < 0.001); **F**) Boxplot comparing distance of CD31/CD34- positive endothelial cells of Ki67-high and Ki67-low levels from Neurofilament high endothelial cells (log2 transformed; p < 0.001). P-values are based on Mann-Whitney U test.

To elucidate whether markers of neural units and vasculature are closely localized, we used density estimation of the Neurofilament (NF) marker from low to high expression (**Figure 1c**). Vascular proliferation was determined by CD31CD34 expressing endothelial cells based on high or low Ki67 expression using density estimates from the flowDensity software (19), resulting in two subsets: Ki67-high/CD31CD34+ and Ki67-low/CD31CD34+. We observed that proliferating vessels (Ki67-high/CD31CD34) were clearly distinct in the 2D space from non-proliferating endothelial cells (Ki-67-low/CD31CD34+) (**Figure 1d**).

Comparing the overall distribution of NF expressing cell units associated with endothelial cell subsets (Ki67-high/CD31CD34+ and Ki67-low/CD31CD34+) indicated that NF expression was on average higher in association with proliferative endothelial cell units (Ki67-high/CD31CD34+) than in Ki67-low/CD31CD34+ (P<0.001) (**Figure 1e**). Distance analysis indicated that NF-high cells were significantly closer to proliferative endothelium (**Figure 1f)** (P<0.001) suggesting an association of neurogenic and angiogenic structures and the potential presence of tumor associated neuro-vascular niches.

Taken together, our spatial single-cell analysis revealed that markers reflecting neurogenesis (NF+) and angiogenesis (Ki67-high/CD31CD34+) are co-localized, supporting a relationship which might hold functional importance. To assess the potential clinical relevance of our findings, we further mapped neural and vascular structures and signatures at the case-level in breast cancer using immuno- histochemistry (IHC) and omics data.

### Neural and vascular tissue markers in primary and metastatic breast cancer associate with aggressive tumor features

For case-based validation of neural and vascular structures in breast cancer, we applied IHC staining for NF and Factor-VIII on tissues from 534 primary cases and 95 matched lymph node metastases.

Among 483 primary tumors with sufficient tissue for examination (90%), microaxons were detected in 102 (21%) cases (**Figure 2a**) and nerve bundles in 126 (26%) cases (**Figure 2b**). We found a median MAD (microaxon density) of 0.058 per mm^2^ and median NBD (nerve bundle density) of 0.080 per mm^2^. Presence of microaxons associated with features of aggressive BC (**Table 1**), whereas no associations were observed between expected pre-existing nerve bundles and primary tumor features. Further, in lymph node (LN) metastases (n=58, after exclusion of micrometastases and cases with insufficient material), microaxons were present in 36 (62%) of metastatic lymph nodes with MAD exhibiting reduced values compared to matched primary tumors (P*=*0.001). Microaxons in LN metastases associated with features of aggressive BC; (**Supplementary Table 2**); nerve bundles were not detected in lymph nodes with metastases.

**Figure 2.**
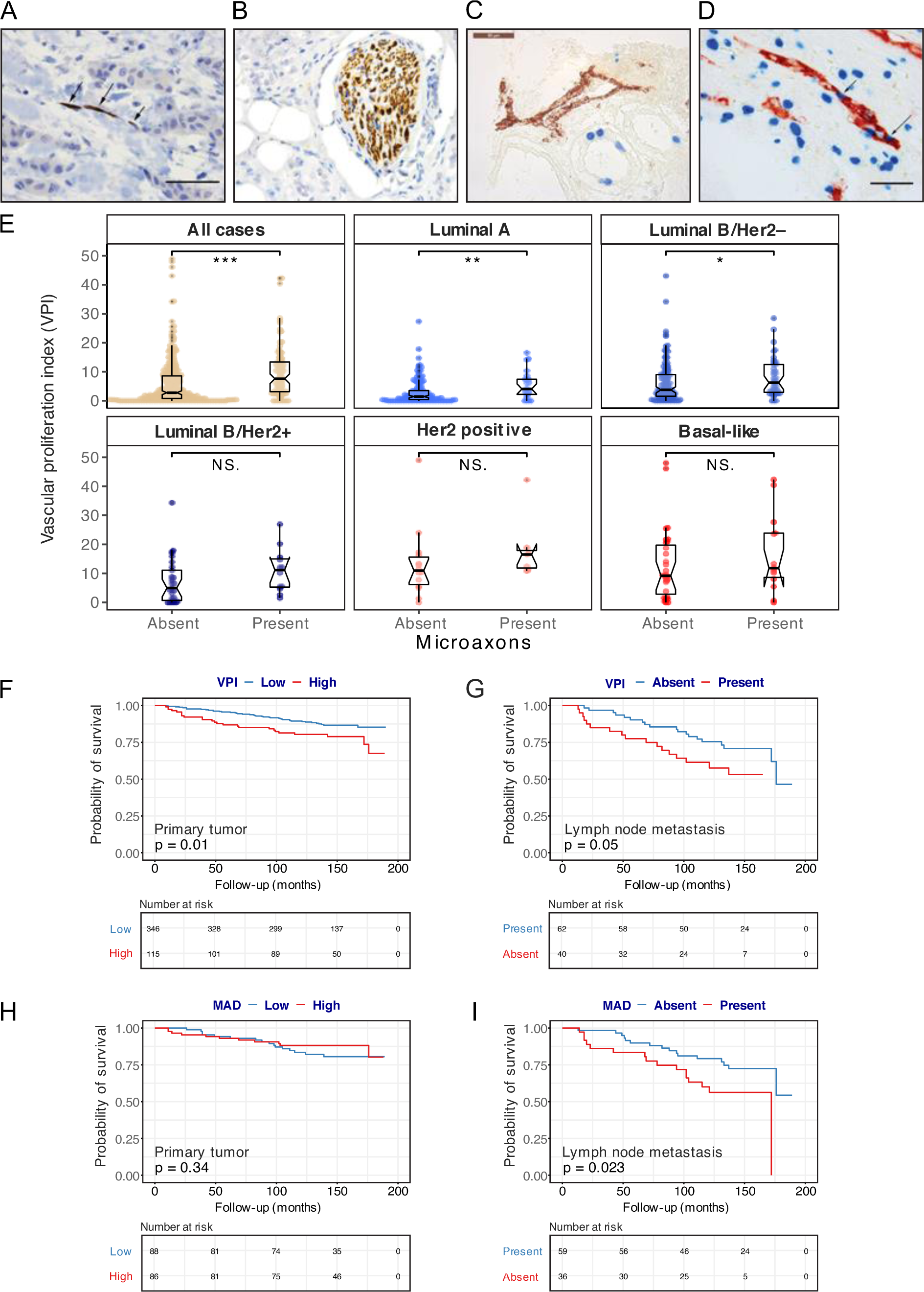
**Immunostaining for markers of angiogenesis and neurogenesis and their associations in primary breast cancer (Bergen Breast Cancer cohort, n=461-483). A**) Immunostaining with Neurofilament antibody (x400), illustrating the positivity in the microaxons between the cords and glands of tumor tissue; **B**) Presence of thick Neurofilament positive nerve bundles; **C-D**) Dual immunostaining with Factor VIII (red) for endothelial cells and Ki67 (blue) for proliferating cells (x400). Vessels with no proliferative activity in (C), and Ki67 positive proliferating endothelial cells in (D) (highlighted with arrows); **E**) Boxplots showing the Vascular proliferation index (VPI) score across breast cancer cases with or without stromal micoraxons in the primary tumor. **F-G**) High vascular proliferation index (VPI) in the primary tumor, and the presence of proliferating vessels in lymph node metastases associates with reduced cancer specific survival in patients; **H**) Microaxon density (MAD) did not associate with survival in primary tumors; **I**) Presence of microaxons in lymph node metastases associates with reduced breast cancer specific survival. All P values shown in (E) are based on Mann-Whitney U test with ‘*’ <0.05, ‘**’< 0.01 and ‘***’< 0.001), NS indicates non-significant results. A log-rank test was used to calculate the p-values in the Kaplan-Meier plots (F-I).

**Table 1.** Associations between presence of axons and clinico-pathologic features of primary tumors (n=483)

| Variable | Axons (primary tumor) |  | OR | (95% CI) | P<sup>c</sup> |
| --- | --- | --- | --- | --- | --- |
|  | Absent<br>n (%) | Present<br>n (%) |  |  |  |
| **Tumor diameter** |  |  |  |  | 0.001 |
| ≤ 2 cm | 302 (82) | 65 (18) | 1.0 |  |  |
| >2 cm | 79 (68) | 37 (32) | 2.2 | (1.4-3.5) |  |
| **Histologic grade** |  |  |  |  | <0.001 |
| Grade 1 - 2 | 325 (82) | 71 (18) | 1.0 |  |  |
| Grade 3 | 56 (64) | 31 (36) | 2.5 | (1.5-4.2) |  |
| **Lymph node metastasis** |  |  |  |  | NS |
| No | 281 (81) | 68 (19) | 1.0 |  |  |
| Yes | 96 (74) | 33 (26) | 1.4 | (0.9-2.3) |  |
| **ER** |  |  |  |  | 0.027 |
| Positive | 326 (81) | 78 (19) | 1.0 |  |  |
| Negative | 55 (70) | 24 (30) | 1.8 | (1.1-3.1) |  |
| **PR** |  |  |  |  | NS |
| Positive | 270 (81) | 65 (19) | 1.0 |  |  |
| Negative | 111 (75) | 37 (25) | 1.4 | (0.9-2.2) |  |
| **HER2 status** |  |  |  |  | 0.027 |
| Negative | 335 (81) | 81 (19) | 1.0 |  |  |
| Positive | 46 (69) | 21 (31) | 1.9 | (1.1-3.3) |  |
| **Ki-67 %<sup>A</sup>** |  |  |  |  | <0.001 |
| Low | 291 (85) | 53 (15) | 1.0 |  |  |
| High | 90 (65) | 49 (35) | 3.0 | (1.9-4.7) |  |
| **Molecular subtype<sup>B</sup>** |  |  |  |  |  |
| Luminal A | 165 (88) | 23 (12) |  |  |  |
| Luminal B / HER2 - | 138 (76) | 44 (24) |  |  | 0.003 |
| Luminal B / HER2+ | 30 (70) | 13 (30) |  |  | 0.003 |
| HER2 + | 16 (67) | 8 (33) |  |  | 0.006 |
| Triple negative | 32 (70) | 14 (30) |  |  | 0.002 |
| **Basal-like phenotype (CK5/6)** |  |  |  |  | 0.09 |
| Negative | 336 (80) | 84 (20) | 1.0 |  |  |
| Positive | 45 (71) | 18 (29) | 1.6 | (0.9-2.9) |  |

n: number of patients; OR: odds ratio; CI: confidence interval; ER: Estrogen receptor; PR: Progesterone receptor; HER2: Human epidermal growth factor 2; CK5/6: Cytokeratin 5/6.
<sup>A</sup>Cut-off 30% (upper quartile); <sup>B</sup> St. Gallen 2013; <sup>C</sup>P-values by Pearson's chi-square test.
Missing data on lymph node status, n = 5.

We then examined markers of vascular structures (**Figure 2c-d**), *i.e.,* microvessel density (MVD), vascular proliferation (pMVD) and VPI (vascular proliferation index; pMVD/MVD x 100). Among 461 primary tumors, we found a median pMVD of 3.07 per mm^2^, median VPI of 3.5 %, and median MVD of 77.6 per mm^2^. High pMVD and VPI scores showed associations with features of aggressive tumors, and scores were higher in HER2 positive, triple negative and basal-like subtypes (**Table 2**). Results for LN metastases paralleled the primary tumors (**Supplementary Table 3**). Scores of pMVD and VPI reflecting active angiogenesis had higher values in primary tumors as compared to matched LN metastases (**Supplementary Table 4**). Within each molecular subtype, we also found that the VPI scores were higher in primary tumors as compared to matched LN metastases (**Supplementary Table 5**).

**Table 2.**
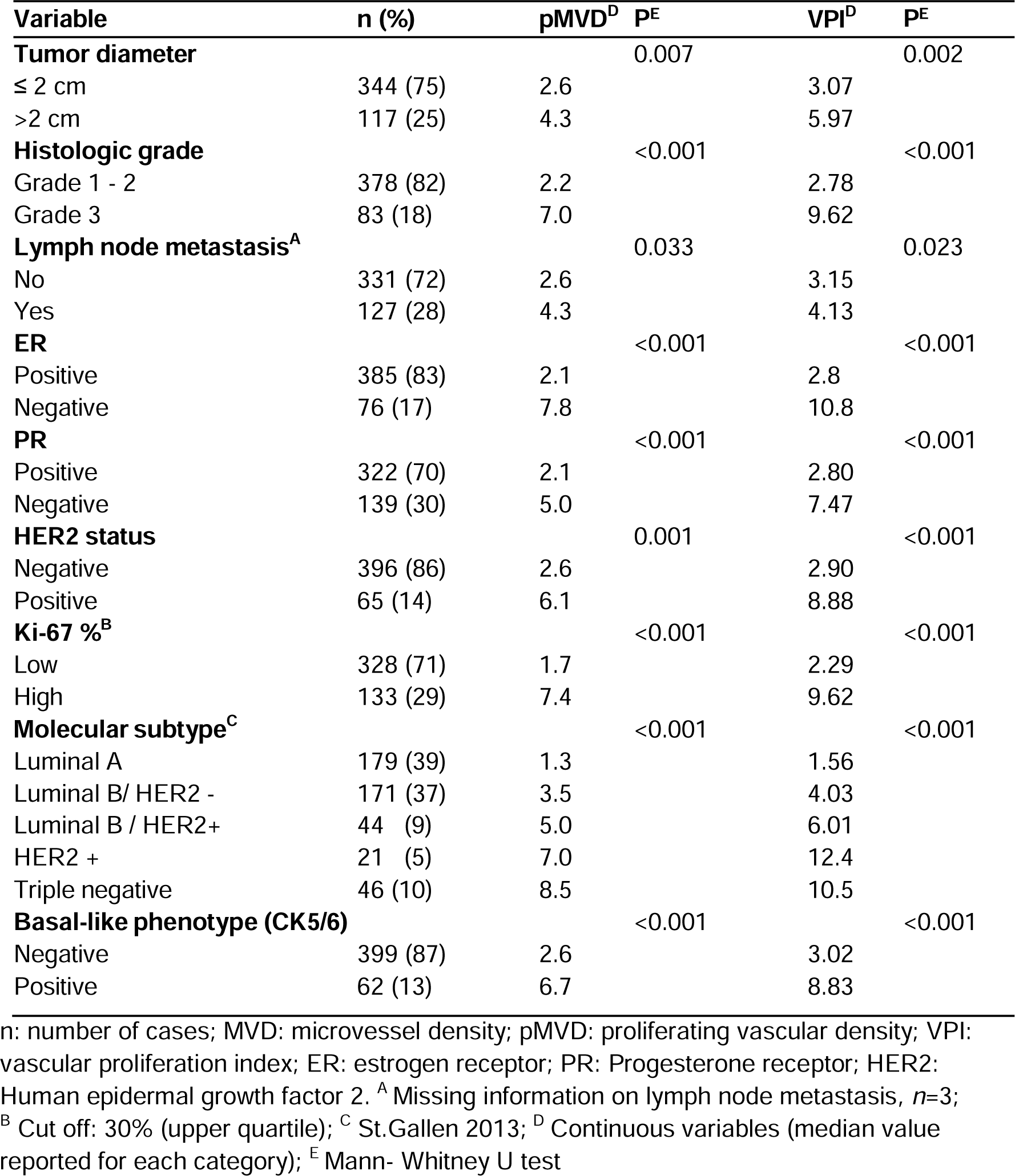
Associations between markers of vascular proliferation (pMVD, VPI) and clinico-pathologic features of primary tumors (n= 461).

| Variable | n (%) | pMVD<sup>D</sup> | P<sup>E</sup> | VPI<sup>D</sup> | P<sup>E</sup> |
| --- | --- | --- | --- | --- | --- |
| **Tumor diameter** |  |  | 0.007 |  | 0.002 |
| ≤ 2 cm | 344 (75) | 2.6 |  | 3.07 |  |
| >2 cm | 117 (25) | 4.3 |  | 5.97 |  |
| **Histologic grade** |  |  | <0.001 |  | <0.001 |
| Grade 1 - 2 | 378 (82) | 2.2 |  | 2.78 |  |
| Grade 3 | 83 (18) | 7.0 |  | 9.62 |  |
| **Lymph node metastasis<sup>A</sup>** |  |  | 0.033 |  | 0.023 |
| No | 331 (72) | 2.6 |  | 3.15 |  |
| Yes | 127 (28) | 4.3 |  | 4.13 |  |
| **ER** |  |  | <0.001 |  | <0.001 |
| Positive | 385 (83) | 2.1 |  | 2.8 |  |
| Negative | 76 (17) | 7.8 |  | 10.8 |  |
| **PR** |  |  | <0.001 |  | <0.001 |
| Positive | 322 (70) | 2.1 |  | 2.80 |  |
| Negative | 139 (30) | 5.0 |  | 7.47 |  |
| **HER2 status** |  |  | 0.001 |  | <0.001 |
| Negative | 396 (86) | 2.6 |  | 2.90 |  |
| Positive | 65 (14) | 6.1 |  | 8.88 |  |
| **Ki-67 %<sup>B</sup>** |  |  | <0.001 |  | <0.001 |
| Low | 328 (71) | 1.7 |  | 2.29 |  |
| High | 133 (29) | 7.4 |  | 9.62 |  |
| **Molecular subtype<sup>C</sup>** |  |  | <0.001 |  | <0.001 |
| Luminal A | 179 (39) | 1.3 |  | 1.56 |  |
| Luminal B/ HER2 - | 171 (37) | 3.5 |  | 4.03 |  |
| Luminal B / HER2+ | 44 (9) | 5.0 |  | 6.01 |  |
| HER2 + | 21 (5) | 7.0 |  | 12.4 |  |
| Triple negative | 46 (10) | 8.5 |  | 10.5 |  |
| **Basal-like phenotype (CK5/6)** |  |  | <0.001 |  | <0.001 |
| Negative | 399 (87) | 2.6 |  | 3.02 |  |
| Positive | 62 (13) | 6.7 |  | 8.83 |  |

n: number of cases; MVD: microvessel density; pMVD: proliferating vascular density; VPI: vascular proliferation index; ER: estrogen receptor; PR: Progesterone receptor; HER2: Human epidermal growth factor 2. <sup>A</sup> Missing information on lymph node metastasis, n=3; <sup>B</sup> Cut off: 30% (upper quartile); <sup>C</sup> St.Gallen 2013; <sup>D</sup> Continuous variables (median value reported for each category); <sup>E</sup> Mann- Whitney U test

Further, we observed that presence of microaxons was positively correlated with pMVD and VPI, while no correlation was detected with MVD scores (**Supplementary** Figure 1a-c). Higher VPI scores were observed in primary tumors with microaxons (P<0.001), when combining all molecular subtypes together as well as for Luminal A and Luminal B/HER2D (**Figure 2e**).

To investigate whether presence of neural and vascular markers associate with patient survival, we performed univariate analysis using the Kaplan-Meier method. Higher vascular proliferation measured by VPI score in primary tumors were associated with shorter survival (**Figure 2f)**, whereas there was no difference in patient survival for MVD score. Increased vascular proliferation measured by VPI in LN metastases indicated reduced survival (p=0.05; **Figure 2g**). Notably, low VPI in primary tumors indicated improved survival within the basal-like category (p=0.05).

When including VPI in multivariate survival analysis using Cox modelling, adjusting for traditional prognostic factors (tumor diameter, histologic grade, lymph node status), VPI maintained independent prognostic value (**Supplementary Table 6**).

MAD measured in the primary tumors did not stratify patients in groups of significantly different survival (**Figure 2h**). However, in LN metastases, we observed that presence of MAD associated with reduced survival (**Figure 2i**).

### Proteomic analysis indicates differential expression of neural and vascular proteins in breast cancer tissues and cells

Using our subcohort of microdissected cases (in-house tissues, tumor epithelial component: 12 basal-like, 12 luminal-like breast cancers), our data indicated 967 differentially enriched proteins when comparing basal-like and luminal-like cases (**Supplementary Table 7; Figure 3a**), of which 26 proteins were involved in *axonogenesis* (n=473; gene-sets GO:0007409); we termed these ‘neurogenesis-related’ proteins. Moreover, 17 angiogenesis-related proteins (n=494; gene-set GO:0001525) were also differentially expressed between basal-like and luminal-like breast cancer cases. The neurogenesis and angiogenesis related proteins were combined into a ‘neurogenesis-angiogenesis signature’ of 43 proteins (NAS-43).

**Figure 3.**
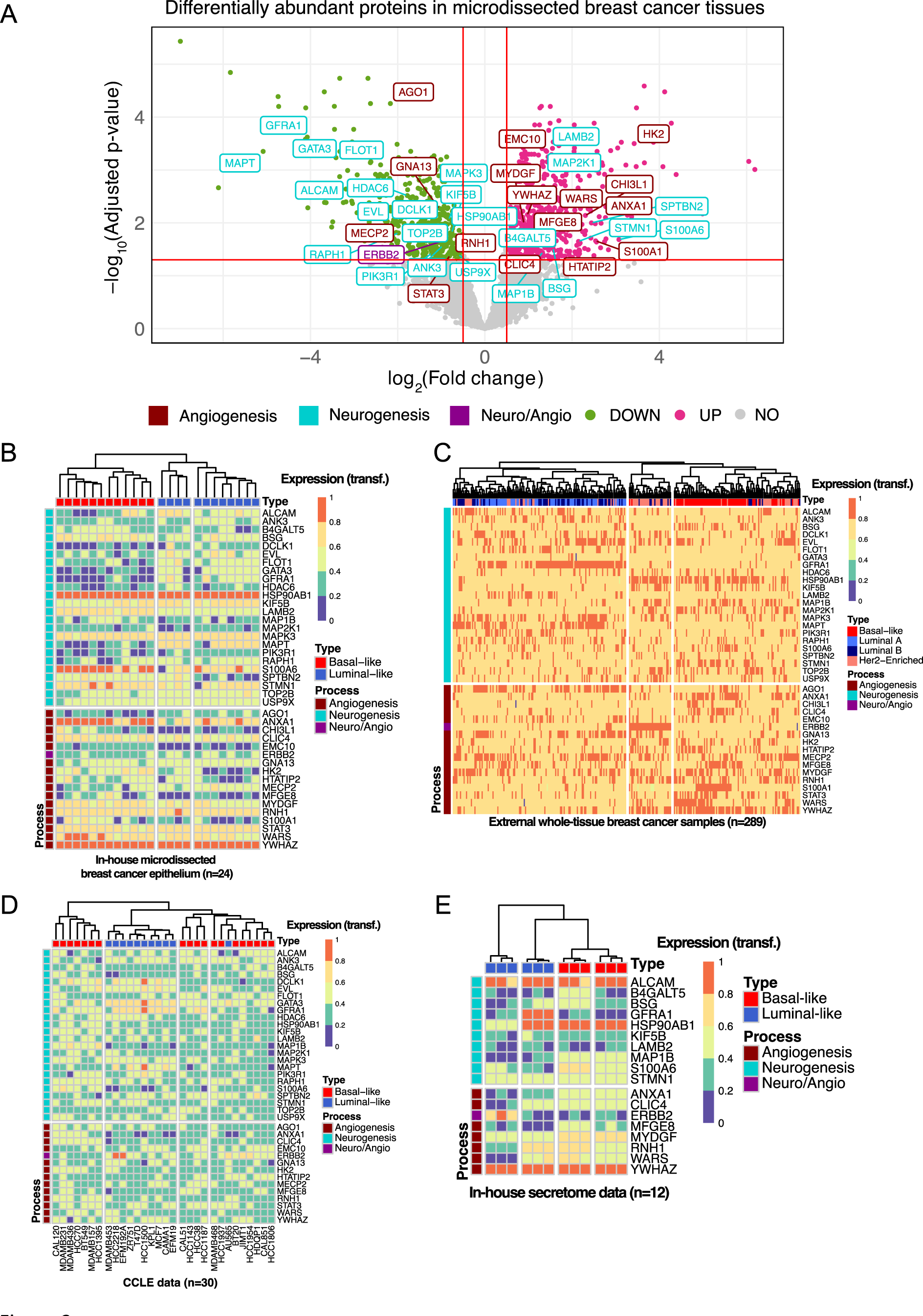
**Differential abundance of neurogenesis-related and angiogenesis-related proteins in breast cancer tissues, cell lines and secretome using mass spectrometry. A**) Volcano plot showing differentially enriched proteins between basal-like and luminal-like tissues. P-values were adjusted using Benjamin-Hochberg method. Down-regulated proteins (green), up-regulated proteins (pink),, angiogenesis-related proteins (dark red), neurogenesis-related proteins (blue), ERBB2 was annotated in both categories (purple); **B)** Heat map presentation of angiogenesis- and neurogenesis-related proteins found in a) using in-house microdissected breast cancer epithelium (n=24); **C)** Publicly-available whole-tissue breast cancer samples (n=289); **D)** CCLE data breast cancer cell lines (n=30); **E)** In-house secretome data (n=12). In all heatmaps the protein expression levels were transformed to an expression scale from 0-1 for comparison purposes.

To probe whether our protein expression profiles could be characteristic for aggressive tumor features, we quantified the NAS-43 proteins using independent datasets from three different sources, in addition to our in-house dataset of microdissected breast cancer cells from which NAS-43 was derived (n=24; **Figure 3b**); 1) a previously reported proteomics profiling of 284 breast cancer cases (17) (**Figure 3c**); 2) a proteomic dataset of 30 BC cell lines from the CCLE (20) (**Figure 3d**), and 3) an in-house secretome dataset derived from four cell lines (**Figure 3e**). Using simple unsupervised hierarchical case clustering, we observed clear separation of basal-like and luminal-like cases (based on Dunn and Silhouette indexes for clustering quality evaluation), see **Supplementary** Figure 2). Taken together, our data indicate that expression profiles of proteins related to neurogenesis and angiogenesis separate basal-like from luminal-like breast cancer subtypes for patient data (microdissected or whole tissues) and cell line information (whole cells or secretomes).

We found the expression of neurogenic and angiogenic proteins, derived from our microdissected breast cancer cohort, to be significantly associated (**Supplementary** Figure 3a). Further, the merged neuro-angiogenic proteomic signature (NAS-43) associated significantly with our NAS signature, a combined neuro-angiogenic mRNA based signature (described below), thus validating a tight association between neurogenic and angiogenic expression profiles (METABRIC cohort; **Supplementary** Figure 2b). Notably, the NAS-43 signature was significantly associated with patient survival as shown in both METABRIC and KMplotter cohorts, in all patients and within the luminal A subgroups (**Figure 4**).

**Figure 4.**
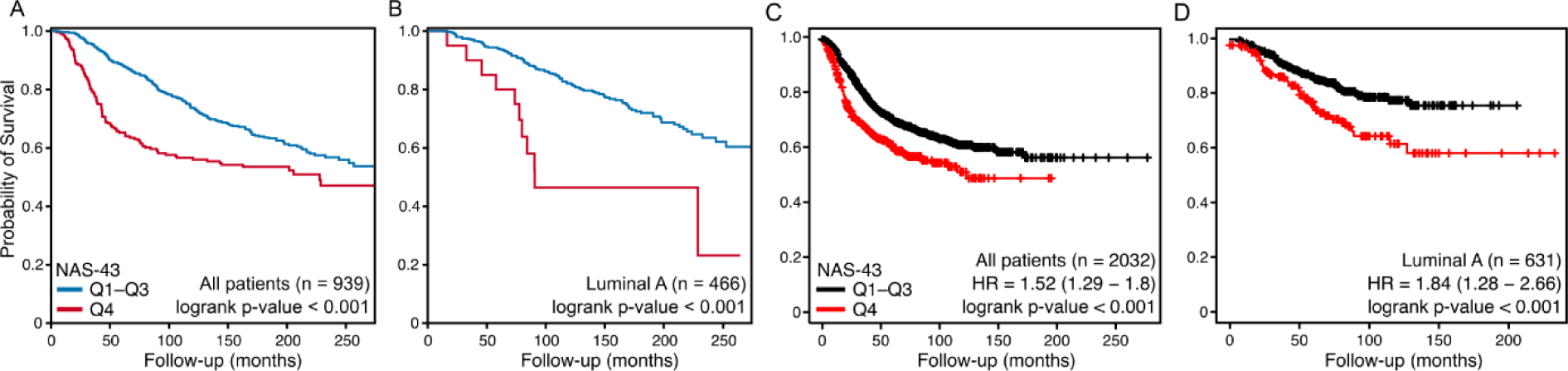
**Univariate survival analysis of the NAS-43 signature in the Metabric Discovery cohort**. Kaplan-Meier plots showing the probability of survival in patients from the METABRIC-Discovery cohort (n = 939; Normal-like subgroup excluded) **(A,B)** and from the online KMPlotter tool **(C,D)** based on their Neuro-Angio gene signature score. Patients with high score (upper quartile) are shown in red and the rest are shown blue (METABRIC) and black (KMPlotter). A high gene signature score is associated with significantly worse survival in all patients **(A,C)** and in patients with Luminal A breast cancer **(B,D)** (log-rank test p ≤ 0.001).

When broadening the focus by assessing neurogenesis and angiogenesis-relevant proteins (n=43) in the independent patient cohort of Asleh et al. (n=284) (17), we found heterogeneity of whole-tissue breast cancer samples. Using simple hierarchical clustering, most of the luminal A cases resided in the same tree branch which was clearly distinct from the basal-like cases. As expected, some luminal B cases and HER2-enriched tumors were clustered together with the main molecular subtypes reflecting the underlying biological characteristics. Our proteomics profiling of neurogenesis and angiogenesis related proteins are in concordance with our IMC and IHC data, providing additional evidence of neurogenic and angiogenic differences between basal-like and luminal-like subtypes.

### Gene expression analysis support neurogenesis and angiogenesis as features of aggressive breast cancer

To further elucidate how gene expression patterns representing neurogenesis and angiogenesis relate to breast cancer phenotypes and tumor progression, we used two previously reported mRNA expression signatures reflecting stroke-related sprouting axons(21) for neurogenesis, and vascular proliferation previously associated with increased microvessel proliferation in endometrial cancer(22) for angiogenesis, examined in the METABRIC cohort (n=1710) combined with breast cancer cell line data from CCLE (n=59) and in-house cell line secretome experiments (12 cell lines; 6 luminal, 6 basal).

On average, sprouting axon score (SAS) and vascular proliferation score (VPS) were higher in basal-like breast cancer as compared to luminal A, luminal B and Her2 enriched subtypes (P<0.001) (**Supplementary** Figure 4a-b). To investigate the combined effect of SAS and VPS, and since these signatures were significantly associated, we merged the scores creating a new Neuro-Angio score (NAS) (see Methods). As expected, NAS showed distinct separation between molecular BC subtypes (**Supplementary** Figure 4c). Considering both METABRIC and CCLE data, we found significant and strong positive correlations between SAS and VPS at the gene expression level (**Supplementary** Figure 4d-e). The highest NAS was seen in basal-like breast cancer (**Supplementary** Figure 4c-f). Adding more information at the global transcriptomic level, these observations are in concordance with our findings from IMC and IHC, and our data support an association between neurogenesis and angiogenesis in aggressive breast cancer subtypes.

We then investigated patient survival using this additional layer of information in the METABRIC data. Patients with high SAS and high VPS were associated with reduced survival (**Supplementary** Figure 4g-h). For the combined NAS, we found associations with shorter survival, both in univariate (**Supplementary** Figure 4i) and multivariate analysis, adjusting for basic factors tumor diameter, histologic grade, and lymph node status (**Supplementary** Figure 4j). When adding PAM50-based molecular subtype to the multivariate model, high NAS independently predicted reduced survival (METABRIC: HR=1.4; 95% CI 1.1-1.7; P=0.001). High NAS maintained independent association with survival within luminal tumors (HR=1.39; 95% CI 1.05-1.84; P=0.022), with a similar trend in non-luminal tumors (P=0.09), when adjusting for tumor diameter, histologic grade, and lymph node status.

Further supporting our finding that NAS was related to aggressive tumor features, associations were found between NAS-score and mRNA-based signatures for EMT, TGFb, stem cell features, and hypoxia, in both METABRIC and TCGA cohorts (**Supplementary** Figures 5-6). When clustering dichotomized signature scores reflecting EMT, stemness, and hypoxia, along with phenotypic clinico-pathologic markers in accordance with high or low NAS, phenotypic subgroups among tumors were evident (four clusters, C1-C4; **Figure 5a-b)**. Two of these clusters (C2, C4) had significant prognostic impact independent of the PAM50 classifier and other basic factors (METABRIC cohort) (**Figure 5c**).

**Figure 5.**
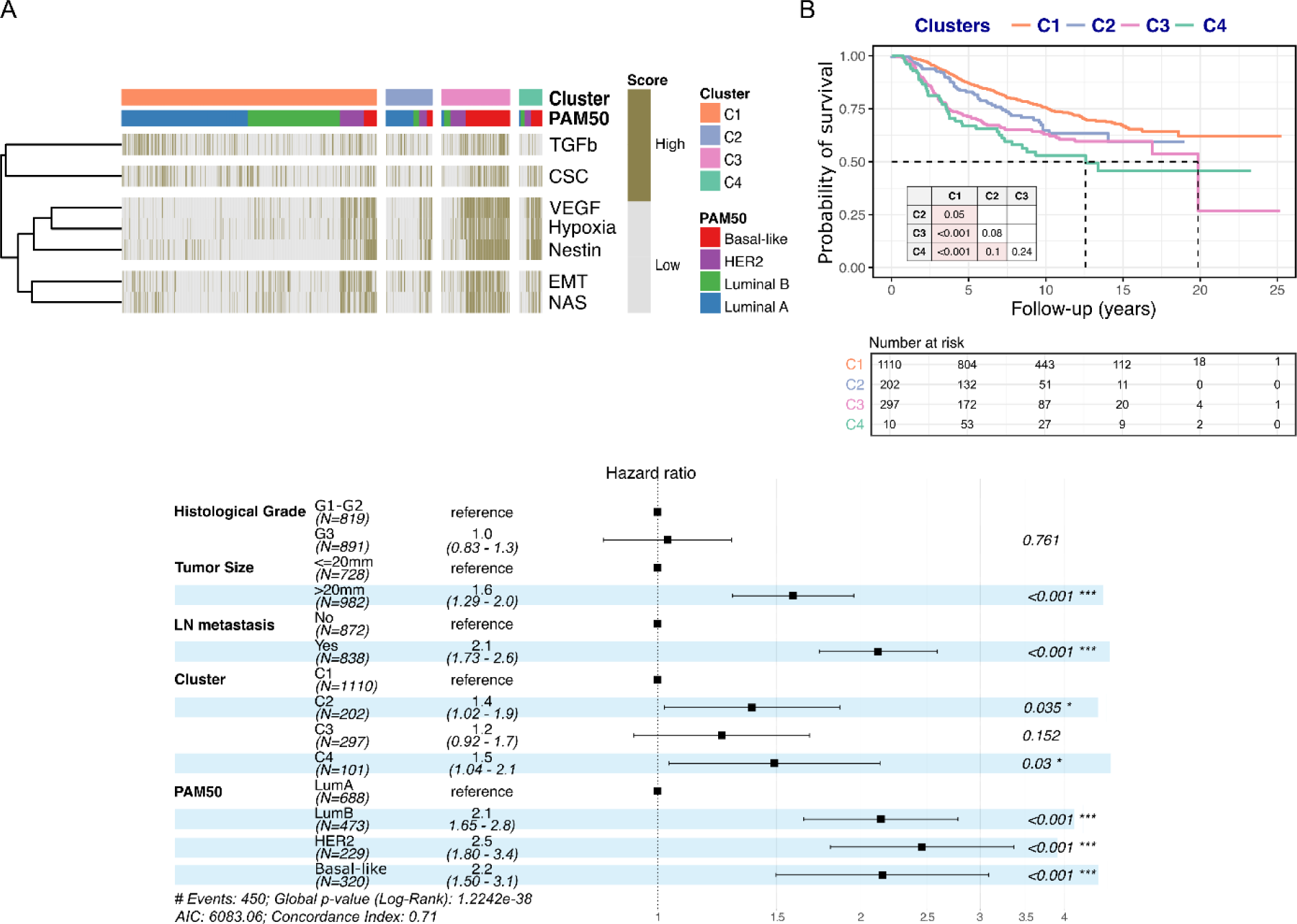
**Unsupervised signature-clustering separates breast cancer patients into high and low breast cancer specific survival**. **A)** Unsupervised Hierarchical clustering of seven breast cancer signatures that are associated with aggressive features revealed four clusters (C1, C2, C3 and C4). The signatures are divided into high and low score. All four breast cancer subtypes are represented in each cluster (C1-C4). **B)** Univariate survival analysis, using the Kaplan-Meier method, shows an association between patient clusters and breast cancer specific survival, with C4 showing the worst survival. **C)** Cox’ proportional hazard modelling shows that the C2 and C4 clusters demonstrate independent prognostic value in multivariate survival analysis, adjusting for tumor size, histologic grade, and LN metastasis and the PAM50 intrinsic subtypes, using data from the combined METABRIC cohort.

## Discussion

The influence of the microenvironment on tumor progress has been increasingly recognized (23–25). Since the link between neurogenesis and angiogenesis has not been well addressed in human cancer, we asked whether these processes are associated traits and related to more aggressive features of breast cancer. By using single-cell tissue analytics in combination with proteomics and transcriptomics data from patients and cell lines, our findings indicate a connection between neural and vascular structures in these tumors, supporting a neuro-vascular niche concept with potential functional implications. Single-cell mapping of breast cancer tissues demonstrated associations between NF expressing cell units and proliferating vessels, and NF+ cells were significantly closer to these proliferating vascular structures compared with the Ki67-low/CD31-CD34+ vessels.

Spatial single-cell mapping of neural and vascular structures indicated significant proximity. Experimental studies have indicated commonalities for neuronal guidance and vascular sprouting, supporting that the two processes might be co-regulated (7, 8), although this has not been previously addressed in human cancer tissues. Future studies will explore a possible functional relationship between neural and vascular components in human breast cancer.

Tumor-associated neurogenesis was strongly associated with markers of high-grade tumors, such as negative hormone receptors, evidence of tumor cell plasticity, and stemness features. The highest level of microaxon density was found in basal-like and Her2 positive breast cancers, while the lowest density was observed in the luminal A subgroup, supporting recent data (15). Nerve bundles, assumed to be pre-existing structures, were observed in many primary tumors but did not associate with any high-grade tumor traits. Such nerve bundles were not observed in lymph node metastases, where microaxons might possibly develop *de novo* from circulating or tissue resident cells (26). Patient survival was reduced in cases with presence of microaxons in lymph node metastases.

At the proteomic and transcriptomic level, combined neuro-angiogenic scores (NAS and NAS-43) was linked to high-grade breast cancer and independently related to reduced patient survival by multivariate analysis. The neuro-angiogenic signature was even stronger than the tumor cell-based classification.

When exploring multiple breast cancer cell lines (27), higher levels of the neuro-angiogenic score were observed in the most aggressive cancer subtypes. This indicates that neuro-angiogenic programs appear to be part of intrinsic tumor cell properties. Taken together, the fact that the neuro-angiogenic score was prognostic within the luminal-like breast cancers, and the finding that some of the cases with high neuro-angiogenic score were hormone receptor positive, might indicate discordant evolution of tumor cells and TME in subsets of breast cancer.

Tumor angiogenesis is an established *hallmark of cancer* (1) and is well described in breast tumors (28, 29). Here, we found consistent and strong associations between proliferating vasculature and aggressive breast cancer phenotypes. Vascular proliferation was increased in basal-like cancers, as we have reported previously(29, 30). As a novel finding, low vascular proliferation within the subgroup of aggressive basal-like cancers indicated a better prognosis compared to the others. Also, elevated vascular proliferation examined in lymph node metastases was associated with reduced survival, which has not been previously reported.

In summary, our data support that neurogenesis and angiogenesis are associated features of aggressive breast cancer. High neuro-angiogenic score was consistently linked to high-grade breast cancer phenotypes and reduced patient survival. Notably, among some hormone receptor positive cases, a high-grade subgroup could be defined by increased neuro-angiogenic score. Single-cell tissue mapping indicated close proximity of neural and proliferating vascular structures, suggesting the possibility of neuro-vascular niches in the tumor microenvironment. Taken together, our findings might be relevant for improved patient stratification and exploration of novel therapy.

## Author contribution

LAA conceived and designed the study and contributed to histologic analyses (IHC, IMC), data interpretation and in writing the major parts of the manuscript. EW contributed to the study design, analyzing the transcriptomic data, general data analyses and interpretation, and in writing the major parts of the manuscript. SA contributed to the study design, performed the tissue-based work and IHC, performed statistical analysis, data interpretation and contributed to writing the manuscript. DK contributed to study design and analyzing the mRNA and IMC data, and contributed to writing the manuscript. HV and KF contributed to analyses of proteomics and IMC data. CA contributed to histologic analyses and work on IMC data, and to collection of clinico-pathologic data. GK participated in data collection and interpretation. KK, AAS, EB, SK, and IMS participated in data collection and interpretation. BD participated in collection of clinical data. All authors commented on and approved the final manuscript. EW and SA made the initial core findings, whereas EW played a larger role in manuscript completion; thus, EW is listed first in the authorship. All authors have read and approved the submitted paper.

## Acknowledgements and funding information

We thank Gerd Lillian Hallseth and Bendik Nordanger for their excellent technical assistance. This work was partly supported by the Research Council of Norway through its Centre of Excellence funding scheme, project number 223250. The study was also supported by the Norwegian Cancer Society and Regional Health Trust Western Norway (Helse Vest).

## Data availability statement

In-house proteomics and transcriptomics data will be made available through online repositories (PRIDE and Gene Expression Omnibus, respectively), and R scripts will be available through GitHub, at the time of publication. Clinical data from patients included in this study may be made available to researchers upon request after completion of a Data Transfer Agreement and confirmation of ethical approval.

## Conflicts of interest

The authors declare no competing interests.

## Notes

### Competing Interest Statement

The authors have declared no competing interest.

### Summary of Updates

The revised manuscript includes several new results and has been extensively rewritten. New data includes single cell spatial analyses of neural and vascular markers.

