## Supplementary material for "Single-cell mapping links neurogenesis and angiogenesis in aggressive breast cancer"

Supplementary Methods, pages 2 – 15

Supplementary Figures, pages 16 – 22

Supplementary Tables, pages 23 – 30

### Supplementary Methods

#### *Patient series*

The study cohort included women diagnosed with primary invasive breast cancer as part of the prospective and population-based Norwegian Breast Cancer Screening Program during 1996-2003 (Hordaland County, Norway; 10% of the Norwegian population), age 50–69 years at time of diagnosis (median 59 years). Patients with distant metastatic disease at time of diagnosis (stage IV) were not included; nine patients refused to participate in the study, leaving 546 cases for inclusion (403 screen-detected,143 interval detected). After exclusion of 12 cases due to lack of available tissue, 534 cases were finally included as previously described [1]. The patients received treatment according to the national guidelines at the time [1]. Clinico-pathologic features like tumor size, histologic type, histologic grade, and hormone receptor (ER, PR) and HER2 status were recorded, as previously described [1]. Histologic type was assessed according to WHO criteria, and histologic grade was determined by the Nottingham criteria [2]. Regarding molecular subtypes, the St.Gallen 2013 consensus criteria were used, with some modifications; cut-off point for the positivity of ER and PR was 10% according to national guidelines at the time [3]. The cut-point for ER and PR positivity was kept at 10 % in this research study, despite the ASCO/CAP guidelines[4], since studies have reported that tumors with ER 1-10% have characteristics similar to those with ER <1% [5, 6]. The basal-like phenotype was determined by the positivity for one basal marker (Cytokeratin 5/6; CK5/6). The tumor block that had the largest tumor or most high-grade area was selected for immunohistochemical studies. Follow-up information, including cause of death, was recorded from medical journals, and cross-checked with the Norwegian Cause of Death Registry. The last follow-up date was December 31, 2011. During this period, 79 (15%) patients died of breast cancer, and 62 (12%) died from other causes. The 5-year breast cancer specific mortality was 7% (37/546). The median follow-up time for survivors was 12.5 years (range 8.3-15.8 years).

#### *Imaging mass cytometry (IMC)*

IMC uses metal-labelled antibodies combined with laser ablation and mass-spectrometry to produce high dimensional images which are further segmented into single-cell information to investigate co-expression and spatial distribution of various markers.

*IMC breast cancer cohort*. We established the IMC breast cancer cohort by merging tissues from two independent tissue collection studies. Cohort-1 consisted of 41 tumor tissues sourced from the Norwegian Breast Cancer Screening Program (9 luminal A, 14 luminal B, and 18 triple negative basal-like breast cancers). Cohort-2 was established in collaboration with Prof. William D. Foulkes at McGill University, Canada, and comprised of 76 tumor tissues (17 luminal A, 16 luminal B, 9 Her2-positive and 34 basal-like breast cancers. We assessed both cohorts using tissue microarrays (TMAs) with 1mm cores in triplets and analyzed one core from each of the 117 cases.

*IMC panel.* The antibody panel consisted of 35 metal-conjugated antibodies and two free metals, iridium, which binds double stranded DNA, and ruthenium, which binds tissue structures, seemingly non-specifically [7]. Antibodies against cytokeratin (clones AE1/AE3) and CD31 (clone EPR3094) were purchased pre-conjugated (Fluidigm), while CD34 (clone ICO115; Cell signaling technologies) and neurofilament (clone C28E10; Abcam) were conjugated in-house using the Maxar X8 Multimetal Labeling Kit (Fluidigm). CD31 and CD34 were conjugated to the same metal (151 Eu).

*Antibody validation.* All antibodies were validated by IHC using a test-TMA with positive control tissues for all antibodies included in the panel: tonsil, placenta, hippocampus, cerebellum, autonomic ganglion and peripheral nerve tissue, normal breast tissue, and selected breast carcinomas (ER+/PR+, HER2+, ER-/PR-/HER2- and basal-like breast carcinomas). For some markers, our IHC staining was compared to staining performed by the Human Protein Atlas (proteinatlas.org). A pilot-TMA (five basal-like; five luminal-like) was established to validate and calibrate the experimental protocol.

*IMC staining protocol*. Antibody hybridization was performed as described in the "Imaging Mass Cytometry Staining Protocol for FFPE Sectioned Tissue" (Fluidigm) with a few modifications. In short, the freshly cut TMA slide underwent dewaxing, rehydration and antigen retrieval in a Ventana Ultra Discovery Autostainer (Roche Diagnostics GmbH). The antibody mix was applied to the slide, which was then stored overnight at 4°C in a hydration chamber. After antibody incubation, the slide was first washed with PBS with and without detergent, before incubation first in 0.3 uM Iridium (Ir)-intercalator (Fluidigm), washed, and then in 0.0005% Ruthenium ($RuO_4$)/PBS

(Electron Microscopy Sciences) [7]. Finally, the slides were washed in MaxPar $H_2O$, air-dried, and stored at 4 °C.

*IMC acquisition and data pre-processing.* Data from the IMC cohort (n= 117) were acquired by a Helios time-of-flight mass cytometer (CyTOF) coupled to a Hyperion Imaging System (Fluidigm) and administered using the CyTOF Software (v7.0.8493; Fluidigm). The square inscribed in the circular TMA cores were laser ablated at 200 Hz at a resolution of approximately 1µm$^2$.

The IMC data was processed using the ImcSegmentationPipeline [8]. First, the IMC output (.txt files) were converted to TIFF-images (ImcTools v2.1.7), which were used for pixel classification (Ilastik v1.3.3post2) to generate probability maps of nuclei, cytoplasm/membrane, and background. These were used to segment single cells using Cell Profiler (v4.0.7). A cell was identified by its nucleus, and the cytoplasm was defined as the area extending four pixels from the nuclear boundary. The resulting cell masks were loaded into Histocat (v1.7.6; [9]) and exported as csv-files to be used in down-stream analyses. We note that even with high-quality segmentation, the imaging of tissue segments cannot rule out cases of “overlapping cell units” that do not capture the nucleus of an individual cell. Therefore, in a few cases nuclei-mismatched signal can be assigned to neighboring cells, especially in (cellular) dense areas.

*IMC data normalization and analysis workflow.* For each marker in the antibody panel, the total intensity per cell unit was computed. Values were divided by the cell size and were arcsinh-transformed. The single-cell data per channel was censored at the 99$^{th}$ percentile to remove outliers. Single-cell annotation was performed similar to the methodology presented in [10]. The annotation procedure was implemented in a hierarchical scheme using unsupervised clustering/meta-clustering and prior knowledge of cell type defining markers of our antibody panel. There are two steps: initially, single cells were categorized as Immune and Non-Immune. FlowSOM [11] was used to cluster the data into 120 clusters, that further merged based on cluster profile cosine similarity. After this step, Phenograph [12] was used to refine the annotation and split Immune cells to three groups (B cells, T cells, macrophages), and non-Immune cells to three groups (epithelial cells, endothelial cells, stromal cells). For the clustering approach, data were standardized across markers using standard normalization.

For further stratification of stromal and endothelial cell populations, the antibody intensity values of Ki67 and CD31/CD34 markers were Z-scored, and a cutoff of 0.5 was applied to split the cell populations to high and low subgroups. For visualizations of cell units, high-dimensional single-cell data were mapped to 2D using UMAP method[13]. We applied the UMAP implementation in R, with default parameters and seven initial dimensions namely: expression of Ki67, CD31/CD34, Neurofilament, Nestin, Vimentin, aSMA, CD45, pan-cytokeratin antibodies.

To estimate the distance of Ki67 high/low cells to their closest endothelial cells, we used a k-nearest neighbor approach. For every cell of Ki67 high/low group, we first selected all available k-closest cells based on their physical proximity. From them, we selected the endothelial cells, and we computed the Euclidean distance. The average log2-transformed distance was used for visualizations, and the procedure was repeated for different values of k closest neighbors starting from 3 to 10 to assess reproducibility.

***Immunohistochemistry (IHC)***

Sections from primary tumors and the matched axillary node with the largest metastatic focus (≥ 2 mm) were used for assessment of nerves (Neurofilament antibody). For Neurofilament (NF) staining (pan-nerve marker), the monoclonal mouse antibody Neurofilament (DAKO M0762) was used. Briefly, the sections were de-waxed by xylene and rehydrated by ethanol in different concentrations, followed by heat-induced epitope retrieval using DAKO S1699 (Decloaking Chamber Plus, Biocare Medical) for 20 minutes. After cooling down for about 15-20 minutes, an endogenous enzyme block (DAKO K4007) was used to block endogenous enzymatic activity. The slides were then incubated with the primary monoclonal mouse antibody Neurofilament (DAKO M0762), at 1:20 dilution for 60 minutes, followed by the secondary antibody Envision System-HRP Labelled Polymer Anti-Mouse (DAKO K4007) in 30 minutes. The visualization was performed by diaminobenzidine (DAB) as chromogen for 10 minutes, followed by hematoxylin and eosin (HE) for 3 minutes. Sections from cerebral cortex were used as positive control in each run.

Sections from the same block of the primary tumor and the axillary lymph node with metastases that were used for assessment of tumor-related nerves, were applied for evaluation of angiogenesis markers (Factor-VIII/Ki67 antibodies). Briefly, slides were de-waxed by xylene and rehydrated by ethanol in different concentrations, followed by 20 minutes heat-induced targeted epitope retrieval using DAKO S1699 (Decloaking Chamber Plus, Biocare Medical). The sections were then allowed to cool down for about 20 minutes. To block the enzymatic activity, a dual endogenous block (DAKO S2003) was applied for 8 minutes. A simultaneous dual immunostaining for the pan-endothelial marker Factor-VIII and the cell proliferation marker Ki67 (MIB-1) was applied on slides for 90 minutes using a DAKO Autostainer. For Factor-VIII, the polyclonal rabbit antibody (DAKO A0082) diluted at 1:1600 was applied, while for Ki67, a monoclonal mouse antibody (DAKO 7240) diluted at 1:50 was used. The antibody detection was done by secondary goat anti-mouse antibody (Southern Biotech 1031-04) diluted in EnVision System-HRP Labelled Polymer Anti-Rabbit (K4003) at 1:100 and applied for 30 minutes at room temperature. Visualization was done by AEC (DAKO; applied for 10 minutes) for Factor-VIII and Ferangi Blue Chromogen Kit (Biocare Medical, Concord, CA, USA; applied for 20 minutes) for Ki67. No counter staining was applied. The blood vessels in the adjacent normal mammary or fat tissue stained with Factor-VIII and the proliferating tumor cells stained with Ki67 were used as internal controls for these markers. In lymph nodes, the adjacent capillaries and the proliferating lymphoid cells were used as internal control cells for Factor-VIII and Ki67, respectively. The comparison with HE slides was always done to guide the location of primary tumors and metastatic foci in the breast and affected lymph nodes.

*Microaxon and nerve bundle density.* Microaxon and nerve bundle densities for each case were assessed separately in the whole tumor area using light microscopy (MAD=microaxon density; NBD=nerve bundle density). First, the slides were scanned at low magnification (Leitz dialux 22 EB, x100) to determine the tumor area for counting axons and/or bundles. Since there were overall low nerve density and no obvious hotspot areas in any of the cases, we used visual fields x250 to cover the whole tumor area. Microaxons and bundles were identified as structures strongly positive for Neurofilament monoclonal antibody, located between tumor cells. Microaxons were observed as single, small, and thin structures (Figure 1a), while a nerve bundle was identified as a group of axons (Figure 1b). The number of microaxons or bundles per mm2 were determined.

*Microvessel density (MVD)*. As a measure of angiogenesis, the count of all vessels in ten high power visual fields (HPF x400) was performed as described [14], and reported as counts/mm2. Briefly, sections were scanned at low magnification (x25) to select the area with the highest vascularization ("hot-spot") for each case. Within these areas, microvessels were counted in 10 consecutive microscopic fields (Leitz dilux 22 EB; x400, field size 0.228 $mm^2$), and median as well as mean values/$mm^2$ were calculated. Microvessels included both vessel-like structures with a visible lumen and single endothelial cells or cell clusters **(Figure 1c)**, as defined by Factor-VIII positivity according to Weidner [15]. Notably, the selected tumor areas should consist of at least 50% tumor tissue to be considered in the assessment of microvessel density. Areas with intra-tumoral scarring or necrosis were avoided.

*Proliferating microvessel density (pMVD)*. Using the same ten microscopic fields (x400) that were used for microvessel density counts, vessels containing Ki67 positive endothelial cells were counted. Thus, pMVD represents a separate count of vessels with proliferating endothelial cells and was reported as count/$mm^2$ (**Figure 1d**).

*Vascular proliferation index (VPI)*. For estimation of activated angiogenesis, the proportion (%) of vessels with proliferating endothelial cells of the total number of microvessels counted in ten fields (Vascular proliferation index; VPI) was calculated.

Inter- and intra-observer variability for neurogenesis and vascular markers. For angiogenesis markers, a training set (n=25) with sections of colon cancer that were stained with Factor-VIII/Ki67 were used to compare inter-observer variability with another researcher (K.K.). The final Kappa agreement was 0.5, P = 0.004 for MVD and 0.6, P<0.001 for both pMVD and VPI. Intra-observer variability (S.A.) for the same variables was also tested after one month, with Kappa agreement 0.6, P<0.001, for all the variables. For neurogenesis, inter-observer variability was tested in a subset of cases (n=18) with another observer (E.W.), with good Kappa agreement (Kappa 0.7, P = 0.015 for axon density; Kappa 0.6, P = 0.05 for bundle density). For intra-observer variability, axon and bundle densities was re-counted in all cases after one month (S.A; Kappa 0.7, P<0.001 for both axon and bundle densities).

***In-house proteomic datasets***

*Microdissected breast cancer epithelial tissue.* Twenty-four (12 basal-like; six luminal A; six luminal B) formalin-fixed, paraffin-embedded (FFPE) breast cancer specimens from the Norwegian Breast Cancer Screening Program (described above) were selected. All basal-like samples were also triple negative, and all luminal samples were estrogen- and progesterone receptor positive, and HER2-negative. The luminal B tumors displayed more than 15 % Ki67-positive nuclei (whole section). All were diagnosed as invasive carcinomas (NST; previously ductal carcinomas). Ten micrometers thick FFPE sections were deparaffinized, rehydrated and stained with hematoxylin. Breast cancer tumor epithelium was laser microdissected (PALM MicroBeam, Zeiss) and pressure catapulted into a tube cap (AdhesiveCap 500 opaque, Zeiss). Depending on the available tissue, 0.8 – 1.9 x 107 μm3 were micro-dissected.

Proteins from the microdissected breast cancer tissue were extracted and enzymatically digested with trypsin using the FFPE-FASP protocol as described in detail by Wiśniewski [16]. The samples were then desalted and cleaned using Oasis HLB μElution plates (Waters, Milford, MA, USA). The microdissected samples were analyzed in its entirety on a Q-Exactive HF mass spectrometer (MS; Thermo Fisher Scientific, Waltham, MA, USA) connected to a Dionex Ultimate NCR-3500RS LC system. Samples were dissolved in 2% ACN/0.1% FA and trapped on the pre-column (Dionex, Acclaim PepMap 100, 2 cm x 75 μm i.d, 3 μm C18 beads) in loading buffer (0.1% TFA) at a flowrate of 5 μl/min for 5 minutes, before separation by reverse phase chromatography (PepMap RSLC, 25cm x 75 μm i.d. EASY-spray column, packed with 2μm C18 beads) at a flow of 200 nL/min. Solvent A and B were 0.1% FA (vol/vol) in water and 100% ACN, respectively. The gradient composition was 5% B from 0-5 minutes, which increased linearly to 8 % from 5-5.5 minutes, to 24 % from 5.5-115 minutes to 35 % B from 115-140 minutes and to 90 % B from 140-155 minutes. Washing and conditioning of the column were performed from 155-170 minutes with 90 % B, and reduced to 5% B from 170-180 minutes. The MS instrument was equipped with an EASY-spray ion source (Thermo Fisher Scientific, Waltham, MA, USA) and was operated in data-dependent-acquisition mode. Instrument control was performed using Q-Exactive HF Tune 2.4 and Xcalibur 3.0. MS spectra were acquired in the scan range 375 - 1500 m/z with resolution R = 120,000 at m/z 200, with an automatic gain control (AGC) target of 3e6 and a maximum injection time (IT) of 100ms. The 12 most intense eluting peptides above intensity threshold 5E4, with charge states 2 or larger, were sequentially isolated to a target AGC value of 1e5, with resolution R = 30,000, an IT of 110 ms and a normalized collision energy of 28 %. The isolation window was set to 1.6 m/z with an isolation offset of 0.3 and a dynamic exclusion of 25 seconds. Lockmass internal calibration was used. Raw data from the mass spectrometer was processed using the MaxQuant software (v1.6.0.16) with recommended settings for label-free quantification [17]. Identified features were searched against the human “reference proteome”-database from UniProt.org (downloaded October 2017).

*Secretome from breast cancer cell lines.* We included proteomics secretome data from breast cancer cell lines (n=4), which is previously published by our group [18] (PRIDE unique identifier PXD027136). The dataset consists of secreted proteins from the conditioned media from two luminal-like cell lines (MCF7 and BT-474) and two basal-like cell lines (MDA-MB-231 and Hs 578T).

***In-house transcriptomics dataset***

*Cell lysate from breast cancer cell lines.* Six basal-like and six luminal-like breast cancer cell lines were obtained from the American Type Culture Collection (ATCC, Manassas, VA). Basal-like cell lines: MDA-MB-231 (ATCC© HTB-26™), MDA-MB-468 (ATCC© HTB-132™), SUM159, SUM1315, Hs 578T (ATCC© HTB-126™), BT-549 (ATCC© HTB-122™); Luminal-like cell lines: BT-474 (ATCC© HTB-20™), MCF7 (ATCC© HTB-22™), ZR-75-30 (ATCC© CRL-1504™), T-47D (ATCC© HTB-133), HCC1428 (ATCC© CRL-2327™), SK-BR-3 (ATCC© HTB-30™). All cell lines were cultured media containing 10% Fetal bovine serum (FBS) and 1% Penicillin-Streptomycin (P/S). Additional media supplement: For MB-231 and MB-468 in F12 (Sigma-Aldrich) 1% Glucose, 1% L-glutamine; MCF7 and Hs578T in DMEM (D6429; Sigma-Aldrich), 1% L-glutamine, and human recombinant insulin (Sigma-Aldrich); ZR-75-30, T-47D and BT-474 in RPMI medium with 1%Glucose, and 1% L-glutamine; BT-549 in RPMI-medium with 1% L-glutamine and human recombinant insulin; SUM1315 and SUM159 in F12-medium; HCC1428 in RPMI-medium and 1% L-glutamine; and SK-BR-3 in McCoy’s-medium. Cell cultures were routinely tested for mycoplasma infections (MycoAlert™ Mycoplasma Detection Kit LT07-318; Lonza). Total RNA from the cells was isolated with miRNeasy mini kit according to protocol (Qiagen, Venlo,

NL). Following extraction, RNA quality and yield was assessed using 2100 Bioanalyzer (Agilent Technologies, CA, USA). The global mRNA expression was examined by the Illumina Bead Array Technology (HumanHT-12 v4 Expression Bead Chip; Illumina, CA, USA). For samples passing the quality control (RIN≥7.5), 367 ng of total RNA from each cell line was biotin-labelled and amplified, using the TotalPrep RNA Amplification Kit (Illumina, CA, USA). RNA was reversely transcribed by reverse transcriptase using an oligo (dT) primer bearing a T7 promoter. The resulting full-length cDNA underwent second strand synthesis and clean-up to become a template for in vitro transcription, biotin labelling and amplification with T7 RNA Polymerase. 750 ng cRNA from each sample was then hybridized at 58°C for 19 hours at the Sentrix bead chip microarrays, according to the manufacturer's protocol (Whole–Genome Gene Expression Direct Hybridization Assay Guide; Illumina, CA, USA). Streptavidin-Cy3 was added after hybridization and washing, for signal detection by the Illumina iScan Reader.

Microarray data were feature extracted using Genome Studio Software (Illumina, CA; USA), with default parameters with respect to the control categories [19] as part of the Whole-Genome Gene Expression Direct Hybridization Assay system. . The in-house cell line mRNA gene expression raw data was log2-transformed and further quantile normalized [20]. Potential batch effects were assessed by explorative approaches, using projection and clustering algorithms [21], considering both known biological differences between the cell lines and batch variables.

***Publicly available data sets*.**

For the exploration of gene and protein expression patterns related to neurogenesis and angiogenesis in breast cancer, publicly available microarray mRNA gene expression profiles and proteomic datasets, with information on clinico-pathologic and follow-up data, and molecular subtypes were analyzed. We used the Molecular Taxonomy of Breast Cancer International Consortium (METABRIC) cohorts (discovery cohort, n=939 and validation cohort, n=845)[22], and The Cancer Genome Atlas (TCGA) breast cancer cohort (n=520) [23]. Intrinsic molecular subtypes based on PAM50 classification [24] were available for all cohorts. Normal breast-like cases were excluded from all mRNA data sets. Further, we applied the breast cancer subset of the Cancer Cell Line Encyclopedia (CCLE) data [25] covering 1037 cancer cell lines across different cancer types, including 59 breast cancer cell lines (47 of these with the molecular subtype described) [26] Furthermore, we used data from a recently published proteomics dataset by Asleh and colleagues [27] consisting of 284 patient samples over two cohorts (cohort-1: n=110, collected from 1986 to 1992; cohort-2: n=174, collected from 2008-2013).

***Data analysis, signatures and clustering***

*Proteomic data analysis*. Protein intensities were compared between groups (basal-like and luminal-like) using the Students t-test. Correction for multiple hypothesis testing was done with the Benjamini-Hochberg method [28]. The DEP R package [29] was used for differential analysis of protein expression. For protein selection we used adjusted p-value cutoff of 0.05 and log fold change (logFC) cutoff of 0.5 between the two conditions. Volcano plots of the proteomics results were generated using R package. Gene ontology datasets were collected from the AmiGO 2 resource (version 2.5.12).

*Clustering and visualization of proteomic datasets*. We visualized the expression levels of the identified neurogenesis/angiogenesis-related proteins using heatmaps implemented from the pheatmap R-package. For visualization purposes, in every dataset the protein expression levels were scaled to the interval [0,1]. Hierarchical clustering was performed to the un-scaled values using Euclidean distance and complete linkage. Within the available datasets experimentation was performed to identify the optimized number of clusters. Specifically, hierarchical clustering was repeated multiple times per dataset, and the resulting trees were cut to generate k=2,3,4,5,6,7 and 8 clusters. For each clustering solution we computed two clustering quality metrics namely the Silhouette Score and Dunn index. The clustering solution that maximized both quality metrics was considered the best.

*Gene expression data analyses*. Differentially expressed genes between cases with high versus low neuro-angiogenesis score (cut-off: upper quartile) were identified based on Significance Analysis of Microarrays (SAM) (29). Gene sets significantly enriched in cases with high neuro-angiogenesis score were explored, applying the Gene Set Enrichment Analysis (GSEA; www.broadinstitute.org/gsea) [30], and the Molecular Signatures Database (MSigDB; www.broadinstitute.org/gsea/msigdb). Multiple microarray probes covering the same gene were collapsed according to max probe.

*Sprouting axons and angiogenesis signatures.* Previously published mRNA expression signature reflecting the transcriptional pattern of stroke-related sprouting axons in an experimental model of young mice [31], and a 32-gene signature (vascular proliferation score; VPS) previously associated with increased tissue-based micro-vessel proliferation (pMVD) in endometrial carcinomas [32] were mapped to the METABRIC and TCGA cohorts, and the CCLE data. The sprouting axons score (SAS) was defined as the sum of expression values of the signature genes.

*Angiogenesis signature*. A 32-gene signature (vascular proliferation score; VPS) has previously been associated with increased tissue-based microvessel proliferation (pMVD) in endometrial carcinomas [32]. In this study, VPS was associated with aggressive tumor features and reduced patient survival. 26 of the 32 genes of the original VPS mapped to the METABRIC and TCGA breast cancer data sets. The resulting 26-gene VPS was calculated by subtracting the sum of expression values of down-regulated genes from the sum of up-regulated genes in the vascular proliferation-high cases.

*Consolidated neuro-angiogenesis profile.* A combined neuro-angiogenic score (NAS), was derived by summarizing the sprouting axons and vascular proliferation scores.

*Other gene expression signatures*. For correlation analyses, we mapped two hypoxia signatures [33, 34], an EMT signature [35], a TGF-β signature [36], and two signatures reflecting stemness features [37, 38], to the mRNA data we analyzed. The corresponding signature scores were calculated as described in the primary publications, and if not originally specified, by a sum of expression values of signature genes.

*Overlap between signature gene sets*. We assessed the overlap between gene expression signatures using the RVenn R package. Gene identifiers obtained from the original publications were converted to Entrez identifiers to unify gene names with different aliases. Signatures with overlapping genes were visualized using binary heatmaps (i.e., gene present or absent). The overlap between two signature gene sets was visualized using Venn diagrams. The Jaccard similarity coefficient (Jaccard index) was used to quantify the similarity between the signature gene lists.

*Clustering analysis of signature scores*. Hierarchical clustering (Euclidean distance and complete linkage) was performed using the score values of TGFb, CSC, VEGF, Hypoxia, Nestin, EMT and NAS signatures as input. To increase the data size, for this analysis we used the combined METABRIC cohort (n=1700). The datasets were processed using the Limma R package to correct for batch effects. Experimentation with variable number of clusters was performed to identify the optimized clustering solution by maximizing Silhouette Score and Dunn index. Heatmap visualization of the signature scores was performed using pheatmap R package. For visualization purposes, the signature score values were binarized to high and low based on the 75$^{th}$ percentile.

***Statistics and survival analyses***

Data were analyzed using SPSS (Statistical Package of Social Scienc1es), Version 28.0 (Armonk, NY, USA; IBMM, Corp) and R-packages via Rstudio v1.3.9. A two-sided P-value less than 0.05 was considered statistically significant. Categories were compared using Pearson`s chi-square or Fisher`s exact tests when appropriate. Non-parametric correlations were tested by Spearman's rank correlation, while Mann-Whitney U and Kruskal-Wallis tests were used for comparing continuous variables in groups. Wilcoxon Signed Rank test was used to compare the differences between two continuous variables. The Kolmogorov–Smirnov statistic was used to quantify a distance between the empirical distribution functions of two samples. Odds ratios (OR) and their 95% confidence intervals were calculated by the Mantel-Haenszel method. Kappa statistics were used to test inter- and intra-observer agreement of categorical data. For survival analyses, the endpoint was breast cancer specific survival, defined as the time in months from the date of histologic diagnosis to the date of death of breast cancer. Univariate survival analysis (Kaplan-Meier method) was performed using the log-rank test to compare differences in survival time between categories. Patients who died of other causes or who were alive at last date of follow-up were censored in the analyses. The influence of co-variates on breast cancer specific survival was analyzed by Cox`s proportional hazards method and tested by the backward stepwise likelihood ratio (lratio) test. All variables were tested by log-minus-log plots to determine their ability to be incorporated in multivariate modelling. When categorizing continuous variables, cut-off points were based on median or quartile values, also considering the distribution profile, the size of subgroups, and number of events in survival analyses.

### *Study approval*

This study was approved by the Western Regional Committee for Medical and Health Research Ethics, REC West (REK 2014/1984). Written informed consent was not obtained from the patients, but in accordance with national ethics guidelines and procedures for retrospective studies, all participants were contacted with written information on the study and asked to respond if they objected. In total, 9 patients (1.7%) did not approve of participation.

### *References*


1. Knutsvik G, Stefansson IM, Aziz S, *et al.* Evaluation of Ki67 expression across distinct categories of breast cancer specimens: a population-based study of matched surgical specimens, core needle biopsies and tissue microarrays. PLoS One 2014;9(11):e112121.
2. Elston CW, Ellis IO. Pathological prognostic factors in breast cancer. I. The value of histological grade in breast cancer: experience from a large study with long-term follow-up. Histopathology 1991;19(5):403-10.
3. Goldhirsch A, Wood WC, Coates AS, *et al.* Strategies for subtypes--dealing with the diversity of breast cancer: highlights of the St. Gallen International Expert Consensus on the Primary Therapy of Early Breast Cancer 2011. Ann Oncol 2011;22(8):1736-47.
4. Hammond ME, Hayes DF, Wolff AC, *et al.* American society of clinical oncology/college of american pathologists guideline recommendations for immunohistochemical testing of estrogen and progesterone receptors in breast cancer. J Oncol Pract 2010;6(4):195-7.
5. Fujii T, Kogawa T, Dong W, *et al.* Revisiting the definition of estrogen receptor positivity in HER2-negative primary breast cancer. Ann Oncol 2017;28(10):2420-2428.
6. Iwamoto T, Booser D, Valero V, *et al.* Estrogen receptor (ER) mRNA and ER-related gene expression in breast cancers that are 1% to 10% ER-positive by immunohistochemistry. J Clin Oncol 2012;30(7):729-34.
7. Catena R, Montuenga LM, Bodenmiller B. Ruthenium counterstaining for imaging mass cytometry. J Pathol 2018;244(4):479-484.
8. Zanotelli VRT, Bodenmiller B. ImcSegmentationPipeline: A pixelclassification based multiplexed image segmentation pipeline. In. Zenodo; 2017.
9. Schapiro D, Jackson HW, Raghuraman S, *et al.* histoCAT: analysis of cell phenotypes and interactions in multiplex image cytometry data. Nat Methods 2017;14(9):873-876.
10. Keren L, Bosse M, Marquez D, *et al.* A Structured Tumor-Immune Microenvironment in Triple Negative Breast Cancer Revealed by Multiplexed Ion Beam Imaging. Cell 2018;174(6):1373-1387 e19.
11. Van Gassen S, Callebaut B, Van Helden MJ, *et al.* FlowSOM: Using self-organizing maps for visualization and interpretation of cytometry data. Cytometry A 2015;87(7):636-45.
12. Levine JH, Simonds EF, Bendall SC, *et al.* Data-Driven Phenotypic Dissection of AML Reveals Progenitor-like Cells that Correlate with Prognosis. Cell 2015;162(1):184-97.
13. McInnes L, Healy J, Melville J. Umap: Uniform manifold approximation and projection for dimension reduction. arXiv preprint arXiv:1802.03426 2018.
14. Arnes JB, Stefansson IM, Straume O, *et al.* Vascular proliferation is a prognostic factor in breast cancer. Breast Cancer Res Treat 2012;133(2):501-10.
15. Weidner N, Semple JP, Welch WR, Folkman J. Tumor angiogenesis and metastasis--correlation in invasive breast carcinoma. N Engl J Med 1991;324(1):1-8.
16. Wisniewski JR. Proteomic sample preparation from formalin fixed and paraffin embedded tissue. J Vis Exp 2013; 10.3791/50589(79).

17. Cox J, Hein MY, Luber CA, *et al.* Accurate Proteome-wide Label-free Quantification by Delayed Normalization and Maximal Peptide Ratio Extraction, Termed MaxLFQ. Mol Cell Proteomics 2014;13(9):2513-26.
18. Kjølle S, Finne K, Birkeland E, *et al.* Hypoxia induced responses are reflected in the stromal proteome of breast cancer. Nature Communications 2023;14(1).
19. Illumina. Illumina Technical Note: Gene Expression Microarray Data Quality Control. 2007.
20. Bolstad BM, Irizarry RA, Åstrand M, Speed TP. A comparison of normalization methods for high density oligonucleotide array data based on variance and bias. Bioinformatics 2003;19(2):185-193.
21. Fellenberg K, Hauser NC, Brors B, *et al.* Correspondence analysis applied to microarray data. Proceedings of the National Academy of Sciences of the United States of America 2001;98(19):10781-10786.
22. Curtis C, Shah SP, Chin SF, *et al.* The genomic and transcriptomic architecture of 2,000 breast tumours reveals novel subgroups. Nature 2012;486(7403):346-52.
23. Cancer Genome Atlas N. Comprehensive molecular portraits of human breast tumours. Nature 2012;490(7418):61-70.
24. Parker JS, Mullins M, Cheang MC, *et al.* Supervised risk predictor of breast cancer based on intrinsic subtypes. J Clin Oncol 2009;27(8):1160-7.
25. Ghandi M, Huang FW, Jane-Valbuena J, *et al.* Next-generation characterization of the Cancer Cell Line Encyclopedia. Nature 2019;569(7757):503-508.
26. Barretina J, Caponigro G, Stransky N, *et al.* The Cancer Cell Line Encyclopedia enables predictive modelling of anticancer drug sensitivity. Nature 2012;483(7391):603-7.
27. Asleh K, Negri GL, Spencer Miko SE, *et al.* Proteomic analysis of archival breast cancer clinical specimens identifies biological subtypes with distinct survival outcomes. Nat Commun 2022;13(1):896.
28. Hochberg Y, Benjamini Y. More powerful procedures for multiple significance testing. Stat Med 1990;9(7):811-8.
29. Huber W, Carey VJ, Gentleman R, *et al.* Orchestrating high-throughput genomic analysis with Bioconductor. Nat Methods 2015;12(2):115-21.
30. Subramanian A, Tamayo P, Mootha VK, *et al.* Gene set enrichment analysis: a knowledge-based approach for interpreting genome-wide expression profiles. Proc Natl Acad Sci U S A 2005;102(43):15545-50.
31. Li S, Overman JJ, Katsman D, *et al.* An age-related sprouting transcriptome provides molecular control of axonal sprouting after stroke. Nat Neurosci 2010;13(12):1496-504.
32. Stefansson IM, Raeder M, Wik E, *et al.* Increased angiogenesis is associated with a 32-gene expression signature and 6p21 amplification in aggressive endometrial cancer. Oncotarget 2015;6(12):10634-45.
33. Eustace A, Mani N, Span PN, *et al.* A 26-gene hypoxia signature predicts benefit from hypoxia-modifying therapy in laryngeal cancer but not bladder cancer. Clin Cancer Res 2013;19(17):4879-88.
34. Hu Z, Fan C, Livasy C, *et al.* A compact VEGF signature associated with distant metastases and poor outcomes. BMC Med 2009;7:9.
35. Jechlinger M, Grunert S, Tamir IH, *et al.* Expression profiling of epithelial plasticity in tumor progression. Oncogene 2003;22(46):7155-69.
36. Padua D, Zhang XH, Wang Q, *et al.* TGFbeta primes breast tumors for lung metastasis seeding through angiopoietin-like 4. Cell 2008;133(1):66-77.
37. Kruger K, Wik E, Knutsvik G, *et al.* Expression of Nestin associates with BRCA1 mutations, a basal-like phenotype and aggressive breast cancer. Sci Rep 2017;7(1):1089.
38. Pece S, Tosoni D, Confalonieri S, *et al.* Biological and molecular heterogeneity of breast cancers correlates with their cancer stem cell content. Cell 2010;140(1):62-73.

**Supplementary Figures**

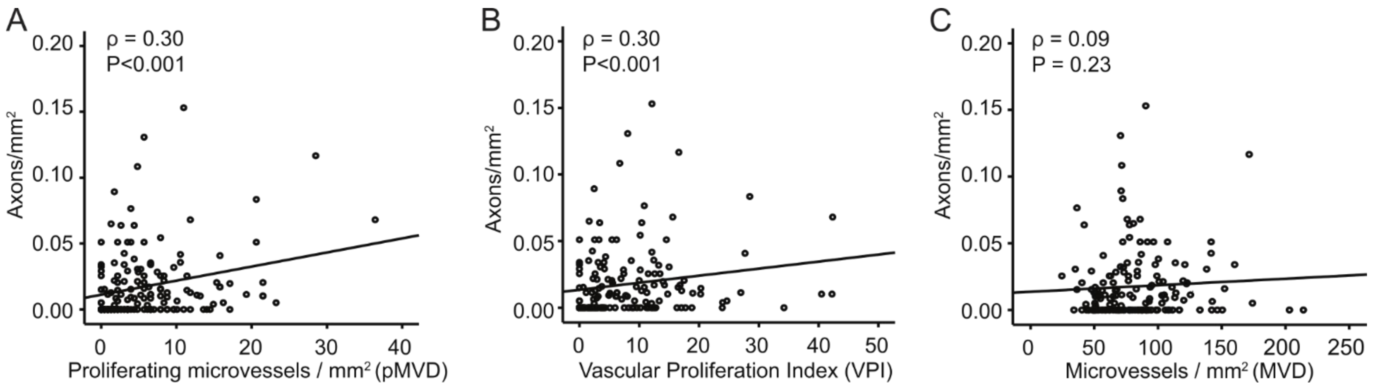


**Supplementary Figure 1**. Correlations analysis between microaxon density and different vascular markers (Bergen cohort; primary tumors). **A**) Scatter plot showing the relationship between microaxon density and proliferative microvessel density (pMVD); **B**) Scatter plot showing the relationship between microaxon density and vascular proliferation index (VPI); **C**) Scatter plot showing the relationship between microaxon density and microvessel density (MVD). The non-parametric correlations were tested by Spearman's rank correlation test.

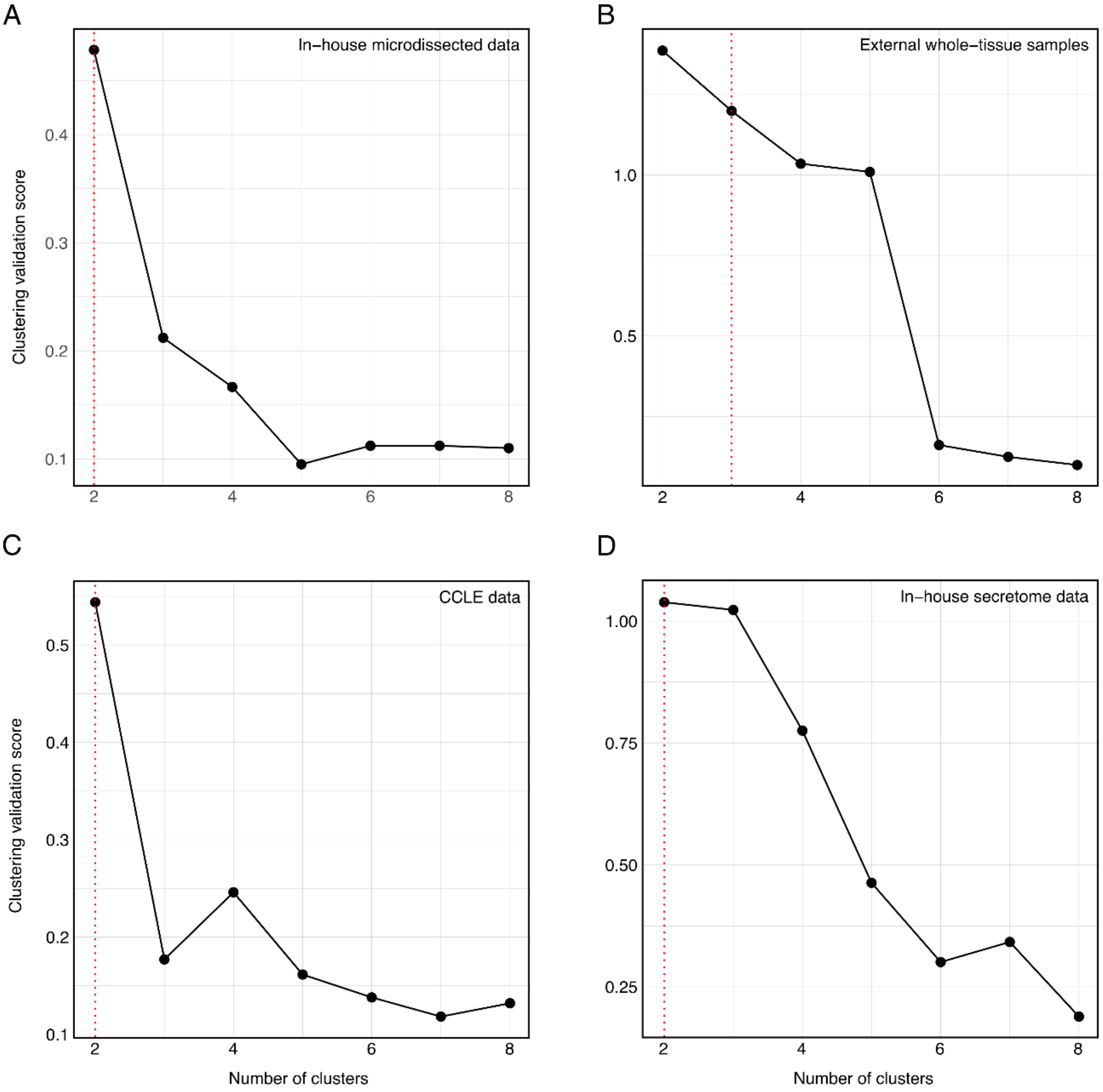


**Supplementary Figure 2. Silhouette Plot for Hierarchical Clustering**. Silhouette plots to evaluate the quality of the clusters identified in Figure 3. A higher clustering validation score (y-axis) means better performance in terms of both the internal cohesion of each cluster and their separation from other clusters. The number of clusters (n) with the highest validation score is considered the "optimal" number of clusters. In all plots (A-D), the "optimal" number of clusters is 2.

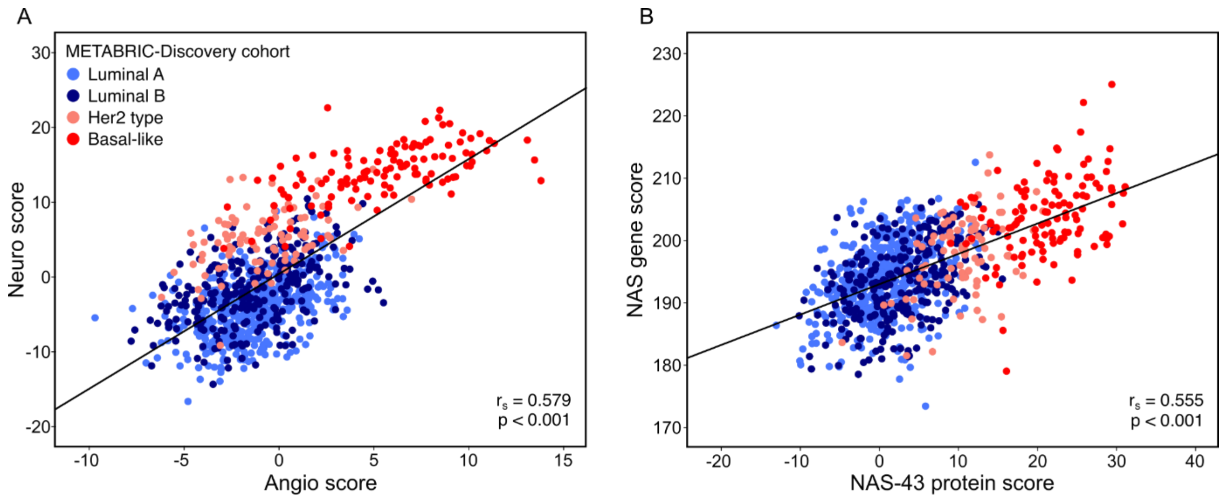


**Supplementary Figure 3**. Scatterplots showing the relationship between **A)** the proteins associated with neurogenesis (n = 26) and angiogenesis (n = 17) in the NAS-43 protein signature (Spearman correlation coefficient = 0.579; $p < 0.001$), and **B)** the NAS-43 protein signature and the previously published NAS gene signature (Spearman correlation coefficient = 0.555; $p < 0.001$). Both scatterplots were made using data from the METABRIC-Discovery cohort.

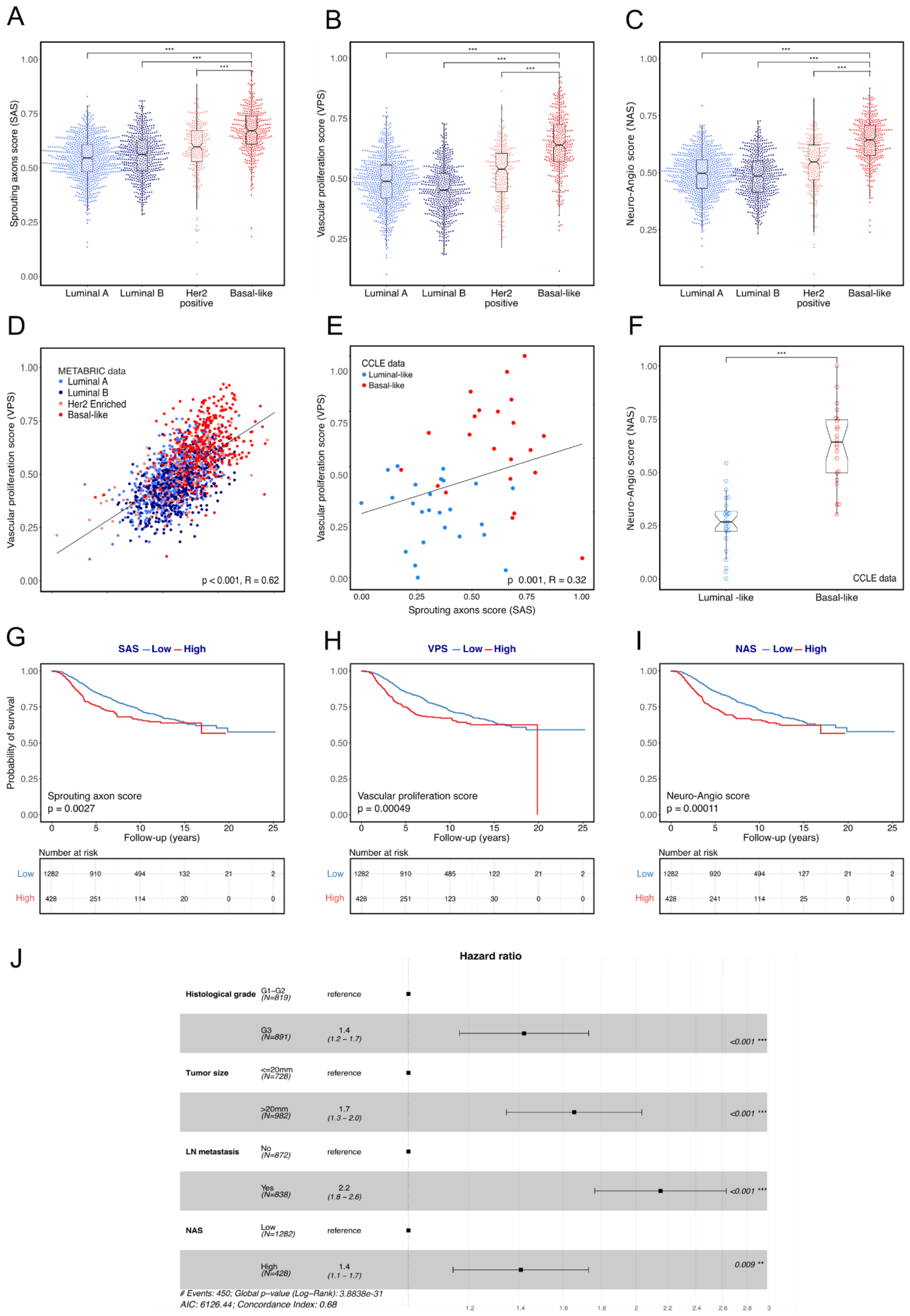


**Supplementary Figure 4. Gene expression analysis of neurogenic and angiogenic features across molecular breast cancer subtypes. A-C)** Bee swarm plots showing the distribution of Sprouting axons score (SAS), Vascular proliferation score (VPS) and the combined Neurogenesis and Angiogenesis score (NAS) across molecular subtypes of breast cancer; **D-E**) Scatter plot showing the relationship between SAS and VPS score values, using data from the METABRIC cohort, and data from all CCLE breast cancer cell lines; **F**) Boxplots showing the distribution of the combined NAS score across molecular subtypes using data from CCLE breast cancer cell lines. **G-I**) High Sprouting axons score (SAS), Vascular proliferation score (VPS), and Neuro-angiogenic score (NAS) associate with reduced breast cancer specific survival in patients from the METABRIC cohort. In all Kaplan-Meier analyses the log-rank test was used to quantify differences. Below every survival curve the number of breast cancer deaths is reported; **J**) Cox' proportional hazard modelling shows that the NAS score demonstrates independent prognostic value in multivariate survival analysis, adjusting for tumor size, histologic grade, and LN metastasis, using data from the combined METABRIC cohort. P values are based on Mann-Whitney U test; '*' <0.05, '**'< 0.01 and '***'< 0.001. Correlations were tested by Spearman's rank correlation test, and the Spearman's correlation coefficient (ρ) is reported. Molecular subtypes are indicated with different colors. In METABRIC data: Luminal A – light blue; Luminal B – dark blue; HER2 enriched – light red; Basal-like - red. In the CCLE data: Luminal-like – blue; Basal-like – red.

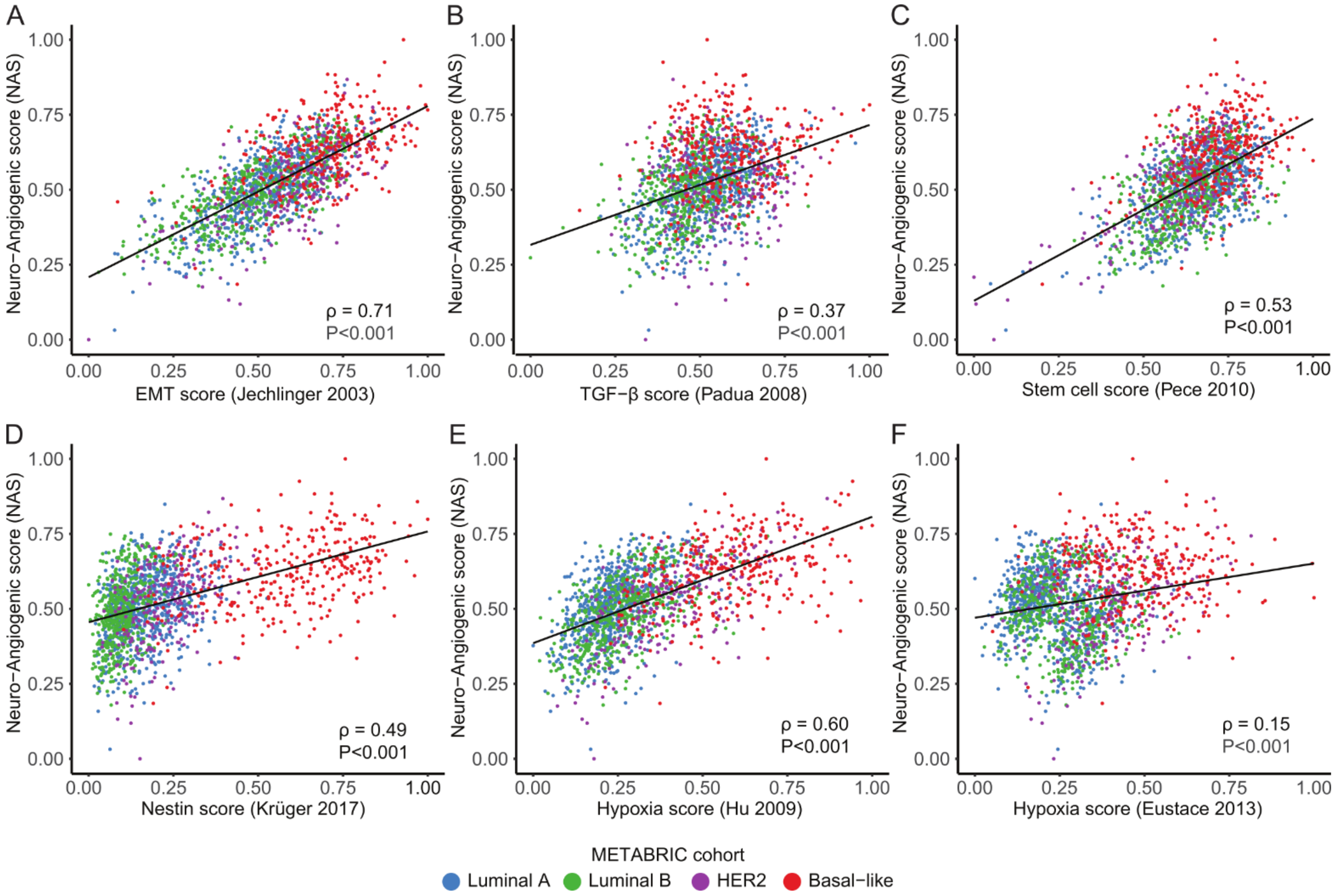


**Supplementary Figure 5**. Investigating associations between Neuro-angiogenic score (NAS) and other transcriptomic signature scores reflecting increased EMT, TGFβ activation, stemness features, and hypoxia in breast cancer, using tissue data from the METABRIC cohort (n=1791). Scatter plots showing the relationship between NAS and scores reflecting: EMT (**A)**, TGFβ activation (**B**), stemness features (**C,D**), and hypoxia (**E,F**). Correlation and coefficients were computed using the Spearman's rank correlation test. Molecular subtypes are indicated by colors: Luminal A - blue; Luminal B – green; HER2 enriched – purple; Basal-like - red.

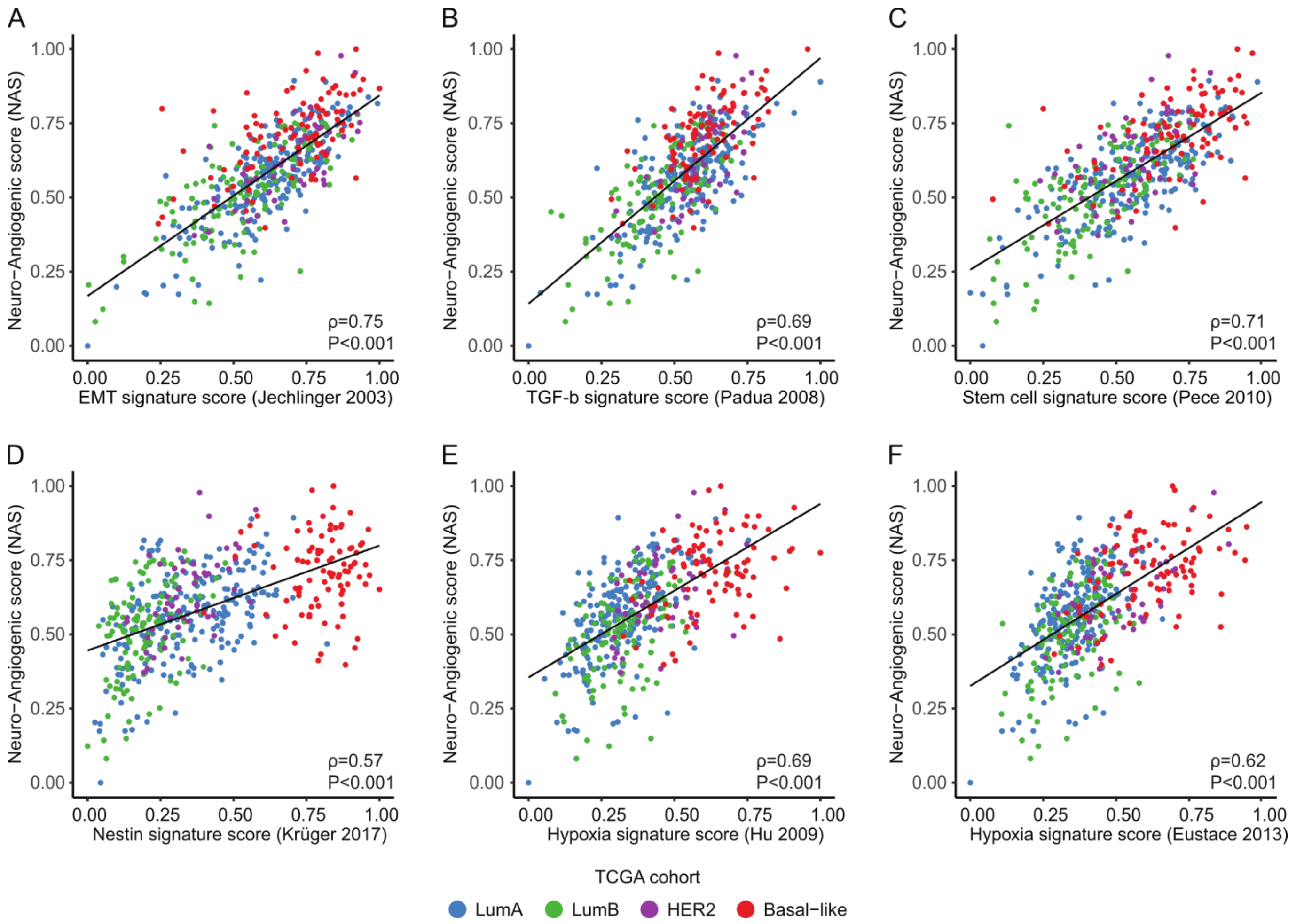


**Supplementary Figure 6**. Investigating associations between Neuro-angiogenic score (NAS) and other transcriptomic signature scores reflecting increased EMT, TGFβ activation, stemness features, and hypoxia in breast cancer, using tissue data from the TCGA cohort (n=505). Scatter plots showing the relationship between NAS and scores reflecting: EMT (**A**), TGFβ activation (**B**), stemness features (**C**, **D**), and hypoxia (**E**, **F**). Correlation and coefficients were computed using the Spearman's rank correlation test. Molecular subtypes are indicated by colors: Luminal A - blue; Luminal B – green; HER2 enriched – purple; Basal-like - red.

### Supplementary Tables

**Supplementary Table 1**: Imaging mass cytometry antibody panel

| Metal isotope | Target | Clone | Dilution | Company |
|---|---|---|---|---|
| 141Pr | α-SMA | 1A4 | 1:100 | Fluidigm |
| 142Nd | Cytokeratin 5/6 | E2T4B | 1:50 | CST |
| 143Nd | Vimentin | D21H3 | 1:100 | Fluidigm |
| 144Nd | p53 | DO-7 | 1:50 | CST |
| 145Nd | PR | SP2 | 1:50 | Abcam |
| 146Nd | Foxa1 | SP88 | 1:50 | CST |
| 147Sm | CD163 | EDHu-1 | 1:50 | Fluidigm |
| 148Nd | CKAE1/AE3 | C11/AE1/AE3 | 1:100 | Fluidigm |
| 149Sm | Stathmin | D1Y5A | 1:4000 | CST |
| 150Nd | PDL-1 | E1L3N® | 1:50 | CST |
| 151 Eu | CD34 | ICO115 | 1:100 | CST |
| 151Eu | CD31 | EPR3094 | 1:25 | Fluidigm |
| 152Sm | CD45 | | 1:100 | Fluidigm |
| 153Eu | CD44 | IM7 | 1:25 | Fluidigm |
| 154Sm | INA | EP676Y | 1:25 | Abcam |
| 155Gd | FOXP3 | D6O8R | 1:25 | CST |
| 156Gd | CD4 | EPR6855 | 1:25 | Fluidigm |
| 158Gd | NCAM | E7X9M | 1:50 | CST |
| 159Tb | CD68 | KP1 | 1:50 | Fluidigm |
| 160Gd | Nestin | 10C2 | 1:50 | Biolegend |
| 161Dy | CD20 | H1 | 1:50 | Fluidigm |
| 162Dy | CD8 | C8/144B | 1:50 | Fluidigm |
| 163Dy | ER | SP1 | 1:25 | Abcam |
| 164Dy | Cytokeratin 14 | SP53 | 1:50 | Abcam |
| 165Ho | PD-1 | EPR4877(2) | 1:25 | Fluidigm |
| 166Er | Her2 | D8F12 | 1:50 | CST |
| 167Er | GATA3 | D13C9 | 1:50 | CST |
| 168Er | Ki67 | B56 | 1:100 | Fluidigm |
| 169Tm | PDPN (D240) | LpMab-12 | 1:50 | CST |
| 170Er | CD3 | Polyclonal | 1:25 | Fluidigm |
| 171Yb | PDGFR-b | 28E1 | 1:25 | CST |
| 172Yb | CTLA-4 | OTI1G10 | 1:25 | Origene/BioNordika |
| 173Yb | DCX | EPR19997 | 1:25 | Abcam |
| 174Yb | Keratin 8/18 | C51 | 1:50 | Fluidigm |
| 175Lu | Neurofilament | C28E10 | 1:200 | Abcam |
| 176Yb | Histone H3 | D1H2 | 1:1000 | Fluidigm |

**Supplementary Table 2.** The associations between the presence of axons in lymph node metastases and clinico-pathologic features in primary tumors (n=95).

| | Axons in lymph node metastases | | | | |
|---|---|---|---|---|---|
| **Primary tumor features** | **Absent n (%)** | **Present n (%)** | **OR** | **(95% CI)** | **P[C]** |
| **Tumor diameter** | | | | | |
| ≤ 2 cm | 38 (70) | 16 (30) | 1.0 | | 0.05 |
| >2 cm | 21 (50) | 20 (50) | 2.2 | (0.9-1.9) | |
| **Histologic grade** | | | | | |
| Grade 1 - 2 | 49 (66) | 25 (34) | 1.0 | | NS |
| Grade 3 | 10 (48) | 11 (52) | 2.1 | (0.8-2.2) | |
| **ER** | | | | | |
| Positive | 52 (67) | 26 (33) | 1.0 | | 0.05 |
| Negative | 7 (41) | 10 (59) | 2.8 | (0.9-3.8) | |
| **PR** | | | | | |
| Positive | 51 (71) | 21 (29) | 1.0 | | 0.002 |
| Negative | 8 (35) | 15 (65) | 4.5 | (1.6-12.3) | |
| **HER2 status** | | | | | |
| Negative | 51 (65) | 27 (35) | 1.0 | | NS |
| Positive | 8 (47) | 9 (53) | 2.1 | (0.8-2.3) | |
| **Ki-67 %[A]** | | | | | |
| Low | 42 (68) | 20 (32) | 1.0 | | NS |
| High | 17 (52) | 16 (48) | 1.9 | (0.8-4.6) | |
| **Molecular subtype[B]** | | | | | |
| Luminal A | 24 (71) | 10 (29) | 1.0 | | |
| Luminal B/ HER2 - | 25 (66) | 13 (34) | 1.2 | (0.4-3.3) | NS |
| Luminal B / HER2+ | 6 (60) | 4 (40) | 1.6 | (0.3-3.9) | NS |
| HER2 + | 2 (29) | 5 (71) | 6 | (0.9-36.2) | 0.03 |
| Triple negative | 2 (33) | 4 (67) | 4.8 | (0.7-30.5) | 0.07 |
| **Basal-like phenotype (CK5/6)** | | | | | |
| Negative | 55 (64) | 31 (36) | 1.0 | | NS |
| Positive | 4 (44) | 5 (56) | 2.2 | (0.3-1.2) | |

ER: estrogen receptor; PR: progesterone receptor; HER2: Human epidermal growth factor 2; OR: odds ratio; CI: confidence interval. [A]cut off 30%; [B]St Gallen 2013; [C]Chi-square test.

**Supplementary Table 3.** Associations between presence of vascular proliferation in lymph node metastases and primary tumor features (n=102).

| | Vascular proliferation in lymph node metastases | | | | |
|---|---|---|---|---|---|
| **Primary tumor features** | **Absent n (%)** | **Present n (%)** | **OR** | **(95% CI)** | **p[C]** |
| **Tumor diameter** | | | | | |
| ≤ 2 cm | 34 (62) | 21 (38) | 1.0 | | NS |
| >2 cm | 28 (60) | 19 (40) | 1.1 | (0.4-2.4) | |
| **Histologic grade** | | | | | |
| Grade 1 - 2 | 52 (66) | 27 (34) | 1.0 | | 0.05 |
| Grade 3 | 43 (44) | 13 (56) | 2.5 | (0.9-6.4) | |
| **ER** | | | | | |
| Positive | 56 (68) | 27 (32) | 1.0 | | 0.004 |
| Negative | 6 (32) | 13 (68) | 4.4 | (1.3-3.5) | |
| **PR** | | | | | |
| Positive | 51 (68) | 24 (32) | 1.0 | | 0.013 |
| Negative | 11 (41) | 16 (59) | 3.0 | (1.2- 2.8) | |
| **HER2 status** | | | | | |
| Negative | 54 (65) | 29 (35) | 1.0 | | 0.065 |
| Positive | 8 (42) | 11 (58) | 2.5 | (0.9-7.0) | |
| **Ki-67 %[A]** | | | | | |
| Low | 44 (69) | 20 (31) | 1.0 | | 0.032 |
| High | 18 (47) | 20 (53) | 2.4 | (1.0-5.5) | |
| **Molecular subtype[B]** | | | | | |
| Luminal A | 25 (71) | 10 (29) | 1.0 | | - |
| Luminal B/ HER2 - | 26 (63) | 15 (37) | 1.4 | (0.5-3.8) | NS |
| Luminal B / HER2+ | 7 (63) | 4 (37) | 1.4 | (0.3-5.9) | NS |
| HER2 + | 1 (13) | 7 (87) | 17 | (1.9-161.1) | 0.007 |
| Triple negative | 3 (43) | 4 (57) | 3.3 | (0.6-17.6) | NS |
| **Basal-like phenotype (CK5/6)** | | | | | |
| CK5/6 - | 56 (61) | 36 (39) | 1.0 | | NS |
| CK5/6 + | 6 (60) | 4 (40) | 1.04 | (0.2-1.9) | |

n: number of patients; OR: odds ratio; CI: confidence interval; ER: Estrogen receptor; PR: Progesterone receptor; HER2: Human epidermal growth factor 2; CK5/6: Cytokeratin 5/6. [A]Cut-off 30% (upper quartile); [B]St. Gallen 2013; [C]P-values by Pearson’s chi-square test.

**Supplementary Table 4.** Angiogenesis markers in matched primary tumors and lymph node metastases (n=127).

| | **Primary tumors** Mean ± SD | **Lymph node metastases** Mean ± SD | **P**[A] |
|---|---|---|---|
| **pMVD** | 6.0 ± 6.8 | 1.1 ± 2.1 | <0.001 |
| **VPI** | 8.1 ± 9.2 | 1.1 ± 2.6 | <0.001 |

pMVD: proliferating microvessel density; VPI: vascular proliferation index; [A]Wilcoxon Rank Sign test

**Supplementary Table 5.** Vascular proliferation index (VPI) in matched primary tumors (PT) and lymph node metastases (LN-mets) across molecular subtypes of breast cancer. Cases with concurrent presence of vascular proliferation in PT and LNmets included in the analyses (n=127).

| | | VPI-PT | VPI-LNmet | $p^B$ |
|---|---|---|---|---|
| **Molecular subtype[A]** | **n** | **Mean ± SD[C]** | **Mean ± SD[C]** | |
| Luminal A | 40 | 4.3 ± 5.6 | 0.5 ± 1.4 | <0.001 |
| Luminal B/ HER2 - | 51 | 7.8 ± 7.8 | 0.8 ± 1.7 | <0.001 |
| Luminal B / HER2+ | 13 | 6.8 ± 7.8 | 0.6 ± 1.0 | 0.01 |
| HER2 + | 12 | 15.0 ± 10.8 | 5.6 ± 5.9 | 0.04 |
| Triple negative | 11 | 17.5 ± 14.9 | 2.2 ± 3.5 | 0.03 |
| **Basal-like phenotype** | | | | |
| CK5/6 + | 17 | 12.1 ± 13.9 | 0.7 ± 1.4 | 0.007 |

n: number of patients; PT, primary tumor; LNmet: lymph node metastases; HER2: Human epidermal growth factor 2; [A]Molecular subtypes, St. Gallen 2013; [B] Wilcoxon Rank Sign test; [C]Mean values are given as median is 0 (many cases showed no vascular proliferation in the lymph node metastases).

**Supplementary Table 6A.** Summary statistics from the differentially enriched neurogenesis-associated proteins between basal-like and luminal- like breast cancer (microdissected tumor epithelial cells)

| Gene | logFC | Average expression | t | P-value | Adjusted P | Direction (B/L) |
|---|---|---|---|---|---|---|
| MAPT | -5.08 | 22.87 | -5.68 | <0.001 | <0.001 | DOWN |
| GFRA1 | -4.07 | 21.36 | -6.18 | <0.001 | <0.001 | DOWN |
| GATA3 | -3.41 | 21.18 | -5.44 | <0.001 | 0.001 | DOWN |
| ALCAM | -3.19 | 23.16 | -4.21 | <0.001 | 0.004 | DOWN |
| FLOT1 | -2.62 | 22.70 | -5.32 | <0.001 | 0.001 | DOWN |
| EVL | -2.09 | 22.80 | -3.75 | <0.001 | 0.009 | DOWN |
| RAPH1 | -1.98 | 22.50 | -3.40 | 0.002 | 0.016 | DOWN |
| DCLK1 | -1.94 | 20.61 | -3.71 | 0.001 | 0.009 | DOWN |
| HDAC6 | -1.78 | 21.06 | -4.37 | <0.001 | 0.003 | DOWN |
| TOP2B | -1.49 | 23.64 | -3.62 | 0.001 | 0.011 | DOWN |
| PIK3R1 | -1.22 | 20.43 | -2.88 | 0.008 | 0.038 | DOWN |
| ANK3 | -0.98 | 22.01 | -3.07 | 0.005 | 0.028 | DOWN |
| ERBB2 | -0.95 | 21.73 | -3.21 | 0.004 | 0.022 | DOWN |
| MAPK3 | -0.82 | 26.16 | -3.77 | <0.001 | 0.008 | DOWN |
| KIF5B | -0.76 | 25.11 | -3.42 | 0.002 | 0.015 | DOWN |
| USP9X | -0.60 | 23.24 | -2.85 | 0.009 | 0.040 | DOWN |
| HSP90AB1 | 0.65 | 30.73 | 3.45 | 0.002 | 0.014 | UP |
| MAP1B | 1.32 | 21.40 | 2.91 | 0.008 | 0.036 | UP |
| MAP2K1 | 1.37 | 20.72 | 4.96 | <0.001 | 0.001 | UP |
| B4GALT5 | 1.46 | 22.01 | 3.23 | 0.003 | 0.021 | UP |
| LAMB2 | 1.55 | 24.77 | 5.79 | <0.001 | <0.001 | UP |
| BSG | 1.57 | 24.01 | 3.01 | 0.006 | 0.031 | UP |
| STMN1 | 2.15 | 25.91 | 3.19 | 0.004 | 0.022 | UP |
| SPTBN2 | 2.44 | 22.54 | 3.67 | 0.001 | 0.010 | UP |
| S100A6 | 3.07 | 27.40 | 3.24 | 0.003 | 0.021 | UP |

**Supplementary Table 6B.** Summary statistics from the differentially enriched angiogenesis-associated proteins between basal-like and luminal- like breast cancer (microdissected tumor epithelial cells)

| Gene | logFC | Average expression | t | P-value | Adjusted P | Direction (B/L) |
|---|---|---|---|---|---|---|
| AGO1 | -2.17 | 20.80 | -7.25 | <0.001 | <0.001 | DOWN |
| MECP2 | -2.01 | 22.68 | -3.69 | 0.001 | 0.010 | DOWN |
| GNA13 | -1.05 | 23.00 | -4.05 | <0.001 | 0.005 | DOWN |
| ERBB2 | -0.95 | 21.73 | -3.21 | 0.004 | 0.022 | DOWN |
| STAT3 | -0.81 | 25.82 | -2.82 | 0.009 | 0.042 | DOWN |
| RNH1 | -0.77 | 26.74 | -3.31 | 0.003 | 0.018 | DOWN |
| MYDGF | 0.89 | 25.22 | 3.67 | 0.001 | 0.010 | UP |
| CLIC4 | 0.93 | 24.91 | 3.03 | 0.006 | 0.030 | UP |
| YWHAZ | 0.98 | 30.20 | 3.87 | 0.001 | 0.007 | UP |
| EMC10 | 1.40 | 20.10 | 5.73 | <0.001 | <0.001 | UP |
| HTATIP2 | 1.93 | 22.04 | 2.85 | 0.009 | 0.040 | UP |
| WARS | 2.06 | 26.38 | 4.00 | <0.001 | 0.006 | UP |
| CHI3L1 | 2.30 | 20.54 | 3.85 | 0.001 | 0.007 | UP |
| MFGE8 | 2.31 | 20.68 | 3.45 | 0.002 | 0.014 | UP |
| S100A1 | 2.54 | 21.34 | 3.21 | 0.004 | 0.022 | UP |
| ANXA1 | 2.76 | 27.67 | 3.80 | 0.001 | 0.008 | UP |
| HK2 | 3.42 | 22.50 | 5.96 | <0.001 | <0.001 | UP |

**Supplementary Table 7.** Multivariate survival analysis (Cox`s proportional hazards method) of vascular proliferation index (VPI) in primary breast cancer compared with standard variables (Bergen Breast Cancer cohort; n=458).

| | | Unadjusted | | | Adjusted | | |
|---|---|---|---|---|---|---|---|
| **Variable** | **n (%)** | **HR[B]** | **(95% CI)** | **P[C]** | **HR** | **(95% CI)** | **P[C]** |
| **Tumor diameter** | | | | | | | |
| ≤ 2 cm | 343 (75) | 1.0 | | | 1.0 | | |
| >2 cm | 115 (25) | 3.6 | (2.2-5.9) | <0.001 | 1.9 | (1.1-3.3) | 0.02 |
| **Histologic grade** | | | | | | | |
| Grade 1-2 | 375 (82) | 1.0 | | | 1.0 | | |
| Grade 3 | 83 (18) | 2.4 | (1.4-4.1) | 0.001 | 1.6 | (0.9-2.8) | 0.12 |
| **Lymph node metastasis** | | | | | | | |
| No | 331 (72) | 1.0 | | | 1.0 | | |
| Yes | 127 (28) | 4.8 | (2.9-7.9) | <0.001 | 3.6 | (2.1 - 6.1) | <0.001 |
| **Vascular proliferation index[A]** | 458 (100) | 1.05 | (1.03-1.07) | <0.001 | 1.03 | (1.01 - 1.06) | 0.04 |

n: number of cases; [A] Continuous variable; [B] Unadjusted HRs given for analyses of 458 cases with data available for all variables in the multivariate analysis;
[C] Likelihood ratio test.
